# Expanding KitBase: Genome and Phenotype Integration of 3,268 Fast-Neutron Rice Mutants

**DOI:** 10.64898/2026.03.31.715674

**Authors:** Artur Teixeira de Araujo, Rashmi Jain, Deling Ruan, Mawsheng Chern, Nguyen Ho, Rohan Bhushan Jhingan, Guotian Li, Phat Q. Duong, Maria Florencia Ercoli, Pamela C. Ronald

## Abstract

Fast-neutron mutagenesis creates diverse genome-wide mutations, providing a powerful tool for crop functional genomics. Here, we present an expanded genomic and phenotypic analysis of 3,268 fast-neutron (FN)-induced mutant rice lines (*Oryza sativa* L. cv. Kitaake). All FN lines were whole-genome sequenced, and mutations were identified by alignment in the Nipponbare and KitaakeX reference genomes. We cataloged over 428,000 mutations affecting 78.49% of Nipponbare genes and 70.38% of KitaakeX genes. *In silico* expression analysis indicates that 575 non-mutated Nipponbare genes are highly expressed and likely essential for viability. Each mutant carries, on average, 68.5 mutations in the Nipponbare alignments or 63.2 mutations for KitaakeX alignments, distributed randomly across all 12 chromosomes with no evident hotspots. FN lines have approximately 8.5% fewer mutations when using the KitaakeX alignment, underscoring the unique contributions of each reference genome and the importance of utilizing both for comprehensive mutation discovery. The majority of mutations are small deletions and single-base substitutions, with deletions predominating in their effect on genes. We found that 74.4% of all transcription factor Nipponbare genes were mutated at least once. Phenotypic characterization of over 2,700 lines revealed a broad spectrum of variation in core agronomic traits (heading date, tiller number, plant height, panicle weight, seed yield components) and other morphological variants of interest. The integration of genomic and phenotypic data through the KitBase platform enabled the identification of candidate genes for several traits of interest. The KitBase website (https://kitbase.ucdavis.edu) has been updated to provide open access to all mutation data and seed stocks, as well as an intuitive query interface, facilitating forward and reverse genetic analyses in rice. This expanded resource enriches the rice functional genomics toolkit and highlights the value of coupling high-density mutation mapping with phenotypic data for rapid gene discovery and crop improvement.

## Introduction

In the face of escalating global challenges such as climate change, food insecurity, and energy shortages, it is imperative to understand the functions of plant genes, which is essential for developing crops with enhanced resilience and productivity, to develop resilient agricultural systems capable of sustaining the growing human population (1,2). Rice (*Oryza sativa*) is one of the world’s most important crops, providing the primary food source for nearly half the global population (3–5). In addition to its critical role in food security, rice serves as an excellent model for monocotyledonous plants, offering significant advantages for functional genomics. Rice has a relatively small genome (∼400 Mb) and extensive genetic resources facilitate genome-wide studies like mutation discovery and mapping (6). Moreover, the wide use of genetic transformation techniques, synteny with other crop species, and a diversified source of related and closely related germplasm further contribute to its utility as a genetic system for functional analyses (7).

Despite its advantages as a model system, a key challenge in elucidating the biological functions of many rice genes is the limited genetic variation within existing cultivated germplasms. This constraint arises from domestication processes and intensive breeding for high-yield traits, which have significantly narrowed the genetic diversity of cultivated rice varieties, making it difficult to associate natural genetic variants with specific phenotypes (8). To overcome this limitation and systematically explore gene function, mutagenesis, which involves inducing artificial genetic variation through chemical or physical mutagens, has been widely employed to establish direct links between specific genetic alterations and their corresponding phenotypic traits (9).

To comprehensively investigate gene function in rice, researchers have developed diverse mutant populations using a range of mutagenesis techniques. These techniques include T-DNA insertions and transposon tagging (e.g., Ac/Ds, Tos17), chemical mutagens (e.g., EMS, MNU), and physical mutagens (e.g., gamma rays, ion beams, and fast-neutrons) (10–14). Each approach contributes with different mutation types and genomic signatures that complement functional genomics pipelines. These efforts have yielded multiple rice mutant libraries in diverse genetic backgrounds (e.g., Nipponbare, Dongjin, Zhonghua 11 and Kitaake) and dedicated databases to facilitate their use (13,15–17). Such community resources have been invaluable, enabling the functional characterization of many rice genes over the past decades.

Fast-neutron (FN) irradiation impacts DNA structure, inducing damage that leads to a spectrum of mutations. While causing single-nucleotide substitutions and small insertions, FN is particularly notable for promoting double-strand breaks that result in larger deletions and chromosomal rearrangements (16). This irradiation approach has been effectively employed to develop mutant populations across various plant species, facilitating functional genomics studies and crop improvement. Notable examples include *A. thaliana* (18), *Hordeum vulgare* (19), *Citrus clementina* (20), *Pisum sativum* (21), *Glycine max* (22) and *O. sativa* (16,23).

In rice, an initial FN mutant population in Kitaake (1,504 M₂ lines) was sequenced at ∼45× coverage, pioneering the concept of a fully whole-genome-sequenced (WGS) mutant library (23). That study identified 91,513 FN-induced mutations affecting 32,307 genes (about 58% of the ∼56k genes in rice). On average, each line carried ∼61 mutations, and the mutation types included single base substitutions (SBS), deletions, insertions, inversions, translocations, and tandem duplications. A high proportion of these mutations were predicted loss-of-function alleles, and in one case, an inversion spanning a single gene was confirmed to cause a short-grain phenotype (23). This work powerfully demonstrated the utility of coupling whole-genome sequencing (WGS) with forward genetics, allowing gene candidates for mutant phenotypes to be pinpointed directly from sequence data without laborious map-based cloning. To share this valuable resource with the community, an open-access database KitBase was established, providing the sequence data and seed stocks for each line. The success and broad utility of this initial Kitaake FN population highlighted the power of next-generation sequencing for genome-wide genotype-phenotype linkage and provided the impetus for further expansion of the resource (23). To broaden the coverage of mutated genes and capture additional mutant information, we further expanded this mutant population and sequenced more lines.

Building on that foundation, the present study significantly expands the KitBase resource by sequencing an additional 1,764 FN mutant lines, bringing the total to 3,268 lines. For the majority of lines, sequencing data was aligned to two reference genomes: the well-annotated Nipponbare reference (IRGSP-1.0) and the newly assembled Kitaake reference genome, KitaakeX (24). This dual reference alignment was motivated by the substantial genetic divergence between Kitaake and Nipponbare, allowing us to maximize mutation discovery and assess how reference choice influences variant calling. In parallel with genomic analysis, we performed extensive phenotypic characterization for over one thousand mutant lines, focusing on key agronomic and developmental traits. By integrating this comprehensive genomic and phenotypic data, accessible through the updated KitBase platform, we illustrate how this resource can accelerate both forward genetics (identifying genes underlying traits of interest) and reverse genetics (finding mutant alleles for genes of interest).

## Materials and Methods

### Plant Materials and Growth Conditions

Rice (*Oryza sativa*) mutant lines were generated and grown as previously described in Li et al. (16), utilizing the parental line X.Kitaake, a *japonica* cv Kitaake carrying the *XA21* gene under the control of the maize ubiquitin promoter. Briefly, 10,000 seeds were mutagenized with 20 grays of FN irradiation, resulting in over 7,300 fertile M1 lines. Sequenced plants were primarily from the M2 generation (Supplemental Data Set 1). Plants for DNA isolation were grown in a greenhouse at the University of California, Davis, under controlled conditions: ∼250 μmol m^−2^s^−1^ light intensity (400-700 nm), 28-30°C, 75-85% humidity, and a 14/10 hour day/night cycle (25).

### DNA Sequencing and Read Mapping

DNA isolation and whole-genome sequencing of the mutant lines were performed according to established protocols (16). Genomic DNA was extracted from 3-week-old plant leaf tissue using the CTAB method (26), quantified with NanoDrop and a fluorometer, and assessed for integrity by agarose gel electrophoresis. Sequencing was conducted on an Illumina HiSeq 2500 platform at the JGI, targeting a minimum 25-fold depth. Reads (2 × 100-bp paired-end) were mapped to the Nipponbare genome version 7 (27) and the KitaakeX genome (24) using the Burrows-Wheeler Aligner-MEM (BWA version 0.7.10) with default parameters (28).

### Genomic Variant Detection

Genomic variant detection largely followed the methodology detailed in Li et al. (16) and Li et al. (23). Samples were processed in groups of up to 50 mutant lines, including a non-irradiated control. A suite of complementary tools, including SAMtools (29), BreakDancer (30), Pindel (31), CNVnator (32), and DELLY (33), was used for variant calling. Variants detected in the parental genome or present in two or more samples within a group were filtered out. SBSs and small Indels (<30 bp) were identified using SAMtools (minimum phred score 100) and Pindel (v0.2.4). Small Indels from Pindel required ≥10 reads, ≥30% variant support, and ≥50 reads in the control line. Large variants (≥30 bp) were called using BreakDancer, Pindel (filtered as above, merging events <10 bp apart), CNVnator (1 kb bin size), and DELLY (for inversions and translocations).

### Functional Annotation of Mutations & Loss-of-Function Mutations

Functional annotation of mutations was performed using SnpEff (34) based on the Nipponbare reference genome version 7 (27) and based on the KitaakeX reference genome (24), as previously described (16). We focused on missense, start/stop codon, and canonical GT/AG splicing site SBSs. Deletions or insertions overlapping exons, as well as inversions or translocations disrupting genic regions, were also included. The gene IDs of transcription factor genes in the rice Nipponbare genome were retrieved from the Plant Transcription Factor Database (PlantTFDB) (35). *In silico* expression analysis was conducted using the Rice RNA-seq Database (https://plantrnadb.com/ricerna/), which compiles expression profiles from 682 RNA-seq datasets spanning 13 distinct tissue types.

### Chromosomal Distribution and Mutation Density Analysis

To evaluate the genome-wide distribution of mutations across the mutant population, mutation density was calculated for each chromosome using a sliding bin approach implemented in R (version 4.3.1) using the packages ggplot2, dplyr, and patchwork. The genome was divided into non-overlapping bins of 500 kb, and each mutation was assigned to all bins it overlapped using an interval overlap criterion, ensuring that mutations spanning multiple bins were accurately represented in each. Mutation density within each bin was expressed as the number of mutations per megabase (mut/Mb), calculated by dividing the mutation count in each bin by the bin size in megabases. Per-chromosome density was computed by dividing the total number of mutations mapped to each chromosome by the chromosome length in megabases, allowing direct comparison of mutation burden across chromosomes of different sizes.

### Cross-Reference of KitaakeX and Nipponbare Gene Identifiers

Conversion between KitaakeX and Nipponbare gene identifiers was performed using two independent resources available through Phytozome (accessed March 2025): (1) a precomputed InParanoid orthology mapping (inparanoid_OsativaKitaake_499_v3.1.tar.gz), and (2) the Best Hit gene correspondence from the KitaakeX annotation file (OsativaKitaake_499_v3.1.P14.annotation_info.txt). A KitaakeX gene was classified as a ’possible ortholog’ of a Nipponbare gene only when both sources independently identified the same gene ID. When multiple ortholog candidates were returned by InParanoid, the top-ranked one-to-one assignment was prioritized. Genes for which no consensus could be established were retained with their original KitaakeX identifier. Full details of these assignments, including the agreement status between the Best Hit and InParanoid sources, are provided in Supplementary Table 3.

### SBS Spectrum and Transition/Transversion Analysis

Single base substitutions were classified using the standard pyrimidine-reference SBS6 convention, in which each of the 12 possible substitution types is collapsed to the corresponding pyrimidine-reference class on the basis of Watson-Crick complementarity (e.g., G>A and C>T represent the same mutational event reported on opposite strands and are combined into the C>T class). SBS6 frequencies were calculated as the proportion of each collapsed class relative to the total number of SBS per dataset. Transition-to-transversion (Ti/Tv) ratios were computed by dividing the combined frequency of C>T and T>C substitutions by the combined frequency of C>A, C>G, T>A, and T>G substitutions. For gene-level spectrum analysis, SBS were intersected with gene body coordinates from the MSU7 annotation (Nipponbare) and the KitaakeX annotation, retaining only mutations overlapping annotated gene features. Nipponbare genes were classified as TE-related or non-TE based on the MSU7 annotation. KitaakeX genes were classified by ortholog transfer using the Nipponbare mapping described in “Cross-Reference of KitaakeX and Nipponbare Gene Identifiers”; genes without a confident ortholog assignment were retained as an “Unmapped” category and analyzed separately to assess potential classification bias. Statistical comparison of SBS6 spectra between gene categories was performed using a chi-square test of independence (6 degrees of freedom), implemented in Python (scipy.stats.chi2_contingency). All analyses were performed in Python 3 using standard libraries (collections, statistics) and visualized using matplotlib.

### Deletion Size Distribution and Bimodality Analysis

Bimodality was assessed using two complementary methods. First, the bimodality coefficient (BC) was calculated as BC = (γ² + 1) / (κ + 3(n−1)²/((n−2)(n−3))), where γ is the skewness and κ is the excess kurtosis of the log-transformed distribution; BC > 0.555 is the standard threshold for detecting non-unimodal distributions. Second, a two-component Gaussian mixture model was fitted to the log-transformed data and compared to a one-component (normal) fit using the Bayesian Information Criterion (BIC); ΔBIC > 10 constitutes strong evidence for two components. Mixture model fitting was performed by splitting the distribution at the trough (log₁₀ ≈ 1.7, corresponding to ∼50 bp) and fitting independent Gaussian distributions to each component. The bimodality coefficient was computed using skewness and kurtosis from *scipy.stats*. A two-component Gaussian Mixture Model was fitted to the log-transformed data using *sklearn.mixture.GaussianMixture* (scikit-learn), and model selection was performed by comparing BIC scores for one- and two-component fits.

### Phenotype Analysis

The phenotypic traits of interest were systematically analyzed to assess plant characteristics. For Germination Rate, twenty to thirty seeds were collected from each plant and allowed to germinate. After seven days, the number of germinated seeds was quantified, and the percentage of germination was calculated. Albino plantlet differentiation frequency was determined by calculating the ratio of albino plants germinated against the total number of germinated plants. Tiller number was recorded as the count of tillers per plant. Days to Heading was measured as the number of days required for the inflorescence to emerge from the flag leaf. Seed number, a key seed yield trait, was quantified by counting the seeds in the first panicle of each plant. Seed Yield was assessed by measuring the panicle weight per plant. Filled grain number was determined by counting the filled grains per panicle. Panicle weight represented the average weight of the panicle from the plants. For panicle area and seed area, samples were placed on a flatbed scanner alongside a ruler as a spatial reference calibration standard. Images were analyzed using ImageJ (https://imagej.nih.gov/), and pixel measurements were converted to cm² using the ruler calibration. Finally, Plant height, a stature and vigor trait, was measured as the height of the whole plant.

For each experimental line, a minimum of three to twenty biological replications were conducted. All measured traits for each line were rigorously compared against a control group grown concurrently under identical conditions. The observed variation within each experimental line was thus attributed to its difference relative to this control. To facilitate the interpretation of these differences, each line’s performance for a given parameter was categorized into specific groups based on its percentage deviation from the control. These classifications are as follows: ’Very High’ for values equal to or above 176% of the control; ’High’ for values between 126% and 175%; ’Normal’ for values between 75% and 125%; ’Low’ for values between 25% and 74%; and ’Very Low’ for values below 24% of the control.

### Identification of Candidate Dwarf and Semi-Dwarf Genes

To compile a reference list of genes associated with dwarf and semi-dwarf phenotypes in rice, we searched five publicly available rice databases using the keywords “dwarf,” “semi-dwarf,” “short stature,” and “compact”: the Rice Annotation Project (RAP; https://rapdb.dna.affrc.go.jp/), Oryzabase (https://shigen.nig.ac.jp/rice/oryzabase/), the Information Commons for Rice (IC4R; https://ngdc.cncb.ac.cn/ic4r/), the Rice Genome Annotation Project (RGAP; https://rice.uga.edu/), and Gramene (https://www.gramene.org/). Redundant entries across databases were manually curated to produce a non-redundant list of candidate genes. The resulting gene list was then queried against KitBase using both Nipponbare (LOC identifiers) and KitaakeX gene identifiers where applicable, to identify FN mutant lines carrying mutations in any of the candidate genes. To confirm the presence or absence of the mutation in *D1/RGA1*, genomic DNA was genotyped by PCR using a single primer set designed to detect both the deletion in FN3664-S and the inversion in FN1535-S (F: 5’-TCTTCACTTAGCACACACAA-3’; R: 5’-TTCCGTTGCTTTGGAACTTT-3’). The wild-type allele produces a 979 bp amplicon; absence of the band indicates homozygosity for the deletion. PCR products were verified by Sanger sequencing to confirm amplification of the expected genomic region.

### KitBase website

The open-access resource, KitBase (http://kitbase.ucdavis.edu/), serves as a comprehensive platform integrating genomic data, mutation information, and seed availability for the Kitaake rice mutant population. Developed using open-source software and tools, KitBase is built upon a MySQL relational database (https://www.mysql.com/) for efficient storage of mutation data. A PHP web interface (http://php.net/) ensures user-friendly data accessibility. Genomic variants aligned to the Nipponbare reference genome are visualized through embedded Variant Call Format (VCF) files within the JBrowse genome browser (36). For sequence-based searches, a standalone BLAST tool (37) has been incorporated. Users can search KitBase using either MSU7 LOC gene IDs (http://rice.plantbiology.msu.edu/) or RAP-DB gene IDs (http://rapdb.dna.affrc.go.jp/). The platform also facilitates seed distribution through a dedicated request webpage. KitBase is hosted by the University of California, Davis.

### Accession Numbers

All sequencing data generated in this study have been deposited into NCBI’s Sequence Read Archive (http://www.ncbi.nlm.nih.gov/sra) under BioProject ID PRJNA385509. Individual line accessions are listed in Supplemental Data Set 1. Sequencing data are also accessible via the JGI website (http://genome.jgi.doe.gov/). Seed stocks for the Kitaake rice mutant lines are available for order through the KitBase platform (https://kitbase.ucdavis.edu/order).

## Results

### Genome Sequencing and Dual-Reference Alignment of 3268 FN lines

To advance functional genomic studies in rice and assess the impact of FN irradiation on the rice genome, we have expanded our established FN-mutagenized population in KitaakeX. The overall strategy for developing this mutant population, suitable for both forward and reverse genetic approaches, is depicted in Figure 1 (16,23). Utilizing Illumina high-throughput sequencing technology, we sequenced an additional 1,764 newly generated FN-mutant KitaakeX lines. All the new lines are M_2_ mutant plants, representing descendants of selfed M1 plants. The mutations in these lines were characterized following the pipeline established by Li et al. (23). To ensure maximal sensitivity and specificity in mutation detection, all newly sequenced lines were aligned against the reference genomes of two *Oryza sativa ssp. japonica* variety: *Nipponbare* (IRGSP-1.0) and *KitaakeX* (a recently published high-quality assembly of *Kitaake*) (24,38). Except for 37 lines, which were analyzed exclusively with the KitaakeX genome.

**Figure 1.**
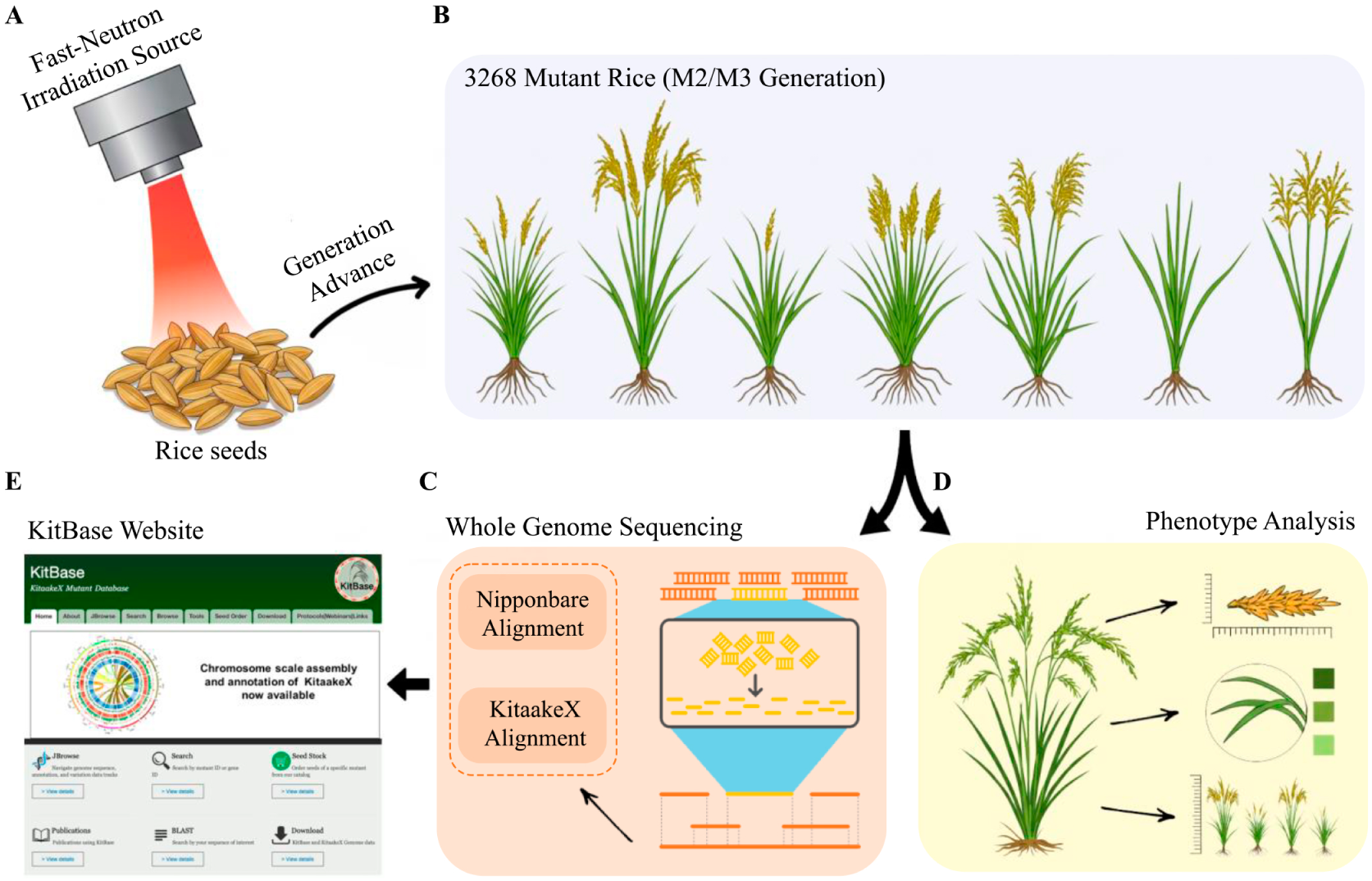
Overview of the strategy employed to develop and expand KitBase, the KitaakeX rice Fast-Neutron (FN) mutagenized population. (**A**) M₀ generation KitaakeX seeds were subjected to FN irradiation, and the resulting M₁ plants were self-fertilized to produce M₂ seeds. (**B**) Seeds from 3,268 M_2_ or M_3_ mutant lines were germinated for genomic DNA isolation and phenotypic characterization. Each line represents an independent mutagenized lineage derived from a single M_1_ plant that was self-fertilized to produce M_2_ progeny. A subset of lines was advanced to the M_3_ generation through an additional round of self-fertilization before analysis. (**C**) Genomic DNA was extracted from leaf tissue of a single M_2_ or M_3_ plant per line. For M_3_ lines, a single individual plant was selected from the M_3_ family for DNA isolation and sequencing, ensuring unambiguous genotype assignment. (**D**) Phenotypic analyses were conducted on a subset of M₂ or M₃ mutant lines. (E) The KitBase website has been updated to include new datasets, analytical tools, and functionalities.

Both reference genome varieties belong to *O. sativa* ssp. *japonica* and are closely related, clustering within the same subpopulation in the 3K Rice Genomes Project classification (38). The Nipponbare genome (MSU v7.0) serves as the gold standard for the rice research community, encompassing 55,986 annotated loci (including transposable element (TE)-related genes) and benefiting from the most comprehensive functional annotation available for rice. In contrast, the KitaakeX genome (35,594 annotated protein-coding genes) enables the analysis of FN lines directly against their cognate reference genome, reducing false positives that can arise from the 331,335 genomic variants (253,295 SNPs and 75,183 InDels) between the two genomes (38). The dual-reference alignment strategy was therefore adopted to leverage the complementary strengths of the two genomes: the KitaakeX reference minimizes false positives arising from natural sequence divergence, while the Nipponbare reference maximizes functional annotation coverage and cross-study comparability within the broader rice research community. A full comparison of the two reference genome assemblies and annotations is provided in Supplementary Table 1.

Furthermore, the 1,504 mutant lines (23), previously analyzed solely with the Nipponbare genome, were reanalyzed using the KitaakeX genome as an additional reference. By combining these reanalyzed lines with the newly sequenced population, KitBase now represents one of the largest fast-neutron-induced rice mutant populations, comprising a total of 3,268 sequenced mutants. This comprehensive dataset includes genetic information from 3,231 lines aligned with the Nipponbare genome and 3,267 lines aligned with the KitaakeX genome (Figure 2-A and Supplementary Table 1), providing a rich resource for the community.

**Figure 2.**
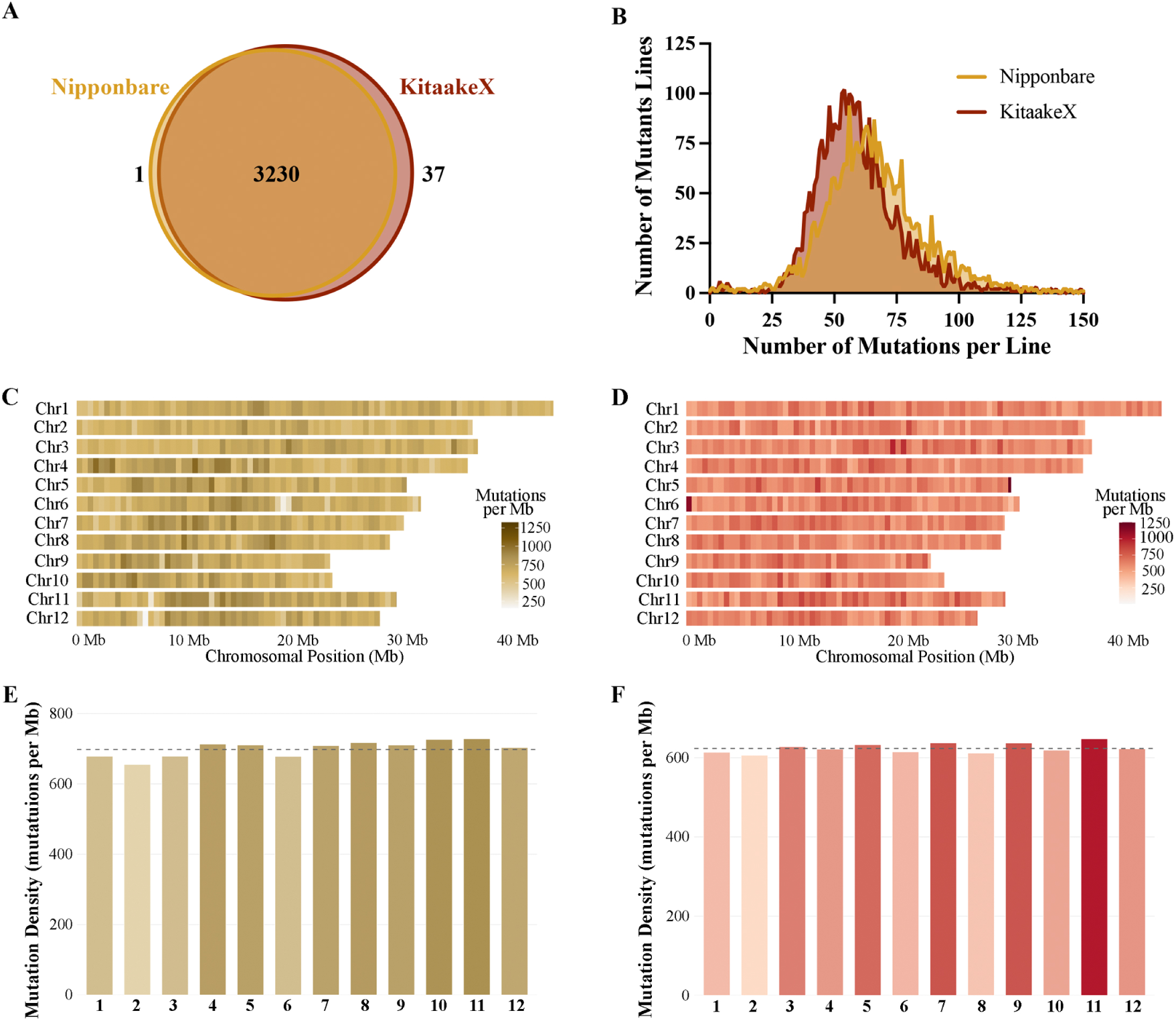
Genome-Wide Characterization of Mutations in 3,268 FN-Induced Rice Mutant Lines. **(A)** A number of mutant lines aligned to each reference genome (Nipponbare and KitaakeX), showing the distribution of sequencing data for comparative genomic analysis. **(B)** Distribution of mutant lines based on the number of detected mutations per line. The data exhibit the distribution until 150 mutations per line. For complete data, refer to Supplementary Table 1. **(C-D)** Chromosomal distribution of mutation density across the **(C)** Nipponbare and **(D)** KitaakeX reference genomes. The genome was divided into 500 kb bins and mutation density (mutations per Mb) is represented as a colour gradient from white (low density) to dark red (high density), capped at 1,250 mut/Mb. Each horizontal strip represents one chromosome. **(E-F)** Per-chromosome mutation density for mutations aligned to the **(E)** Nipponbare and **(F)** KitaakeX reference genomes. Bar height represents the total number of mutations mapped to each chromosome, normalized by chromosome length (mut/Mb). The dashed line indicates the genome-wide average density.

### Mutation Discovery and Reference Genome Comparison

Using the Nipponbare reference genome, a total of 221,481 mutations were identified across 3,231 rice lines (Supplementary Table 1). In parallel, 206,857 mutations were detected in 3,267 lines aligned to the KitaakeX reference. To directly assess the impact of reference choice on mutation detection for individual lines, we compared mutation calls for the subset of 3,230 lines aligned to both reference genomes (Figure 2-A). This analysis revealed a difference of 17,397 more mutations (an 8.52% increase) detected in the Nipponbare alignment compared to KitaakeX. Examining individual lines, we found that the majority (2,335 lines, 72.29%) showed a higher mutation count when aligned to Nipponbare, while only 104 lines (3.22%) had identical counts. Conversely, 24.49% of lines had more mutations called in KitaakeX than Nipponbare.

These discrepancies in mutation counts likely stem from the substantial genomic variations previously reported between the Kitaake and Nipponbare reference genomes, including over 331,000 polymorphisms (24), which significantly impact read mapping and variant calling. While alignments to the KitaakeX genome may reduce false positives by minimizing mapping artifacts caused by sequence divergence, the well-established Nipponbare reference genome offers comprehensive and meticulously curated gene annotation (39). This detailed annotation is fundamental for accurately identifying and understanding the full spectrum of genes affected by mutations. Thus, each reference provides unique advantages in the mutation discovery and interpretation process.

Utilizing both reference genomes provides a more comprehensive mutation catalog for each rice line. Alignments to the KitaakeX genome, which represents the genetic background in which the mutant lines were generated, are expected to reduce false positives due to minimal sequence divergence compared to the mutant lines. Conversely, the Nipponbare genome offers more extensive genomic annotations, facilitating a broader understanding of affected genes. Therefore, results generated from alignments to both references are presented to offer the most comprehensive and robust view of the mutation landscape within the KitBase population.

### Mutation Frequency and Distribution per Line

To further characterize the distribution of mutations across the mutant population, we analyzed the frequency of mutation counts per line using data from both reference alignments (Figure 2-B). The number of mutations per mutant line varied widely but followed an approximately normal (bell-shaped) distribution in the Nipponbare and KitaakeX aligned datasets (Figure 2-B; Supplementary Figure 1-A). This pattern aligns with findings from similar studies (16,23).

The average number of mutations per line is 68.53 in the Nipponbare alignment and 63.15 in the KitaakeX. This relatively low mutation load per line is advantageous for genetic analysis. Specifically, among the 3,268 analyzed lines, the majority harbored fewer than 100 mutations per line (92.88% in Nipponbare and 96.35% in KitaakeX alignments; Figure 2-B). Conversely, highly mutated lines were rare, with fewer than 1.5% exhibiting more than 150 mutations per line. These lines with a higher number of mutations can be valuable for reverse genetics or saturation mutagenesis approaches, as they are more likely to harbor mutations in genes not yet affected in lower-mutation lines. This manageable mutation frequency per line facilitates the identification of causative mutations in downstream genetic screens.

Examining the extremes of the mutation frequency distribution provided further insights. In the Nipponbare alignment, one mutant line (FN-57) showed no detectable mutations (an apparent wild-type), whereas alignment to the KitaakeX genome revealed five mutations in the same line. The lowest number of mutations detectable in the KitaakeX alignment was one, observed in lines FN1110-S and FN422-S (Supplementary Table 1). At the other end of the spectrum, a few lines exhibited significantly higher mutation counts in both alignments: FN3028-S, with 1,042 and 1,123 mutations in Nipponbare and KitaakeX alignments, respectively, and FN3126-S, with 935 and 1,128 mutations. Notably, the KitaakeX alignment identified one additional line with an exceptionally high number of mutations, FN588-S, harboring 4,421 mutations, making it the most highly mutated line in the dataset (Supplementary Table 1). Despite these extremes, the overall mutation distribution remained similar between the two reference genomes (Figure 2-B), indicating that while absolute mutation counts differ slightly, the population-wide pattern of mutagenesis was consistent. We observed no evidence of distinct subpopulations with unusually high or low mutation rates; rather, FN mutagenesis introduced a roughly random number of mutations per line with common central tendencies across the population.

To assess the chromosomal distribution of mutations, all detected FN-induced mutations were mapped and mutation density was calculated across the reference genomes of Nipponbare and KitaakeX (Figure 2:C-F). The analysis revealed an even distribution of mutations across all chromosomes for both alignments, with no evidence of specific chromosomes being more prone to mutations or exhibiting mutational hotspots. In Nipponbare alignment, a genome-wide average density of 698.0 mut/Mb, while the Kitaake-aligned dataset yielded an average density of 623.4 mut/Mb. Comparison of mutation densities across chromosomes confirmed that mutation rates were consistent across the genome, aligning with previous findings in similar mutant populations (23). In both reference alignments, Chr11 exhibited the highest mutation density (728.3 mut/Mb in Nipponbare; 647.6 mut/Mb in Kitaake), while Chr2 showed the lowest (655.0 mut/Mb in Nipponbare; 606.0 mut/Mb in Kitaake). This comprehensive genomic distribution suggests that fast-neutron mutagenesis introduces largely random genomic alterations without strong chromosomal bias, making the resource valuable for probing gene function across the entire genome.

### Affected Genes in 3268 FN-Mutant Lines

Utilizing the established pipeline from Li et al. (23), we identified genes affected by FN-induced mutations, potentially leading to altered gene function, across 3,268 rice mutant lines. Using the comprehensive MSU7 annotation of the Nipponbare genome (27), which includes 55,986 annotated genes (comprising 39,049 non-transposable element (non-TE) genes and 16,937 TE-related genes), we found that 43,946 genes have at least one mutation in the KitBase population. This corresponds to an overall gene coverage of 78.49%, specifically 75.67% (29,550) of non-TE genes and 84.99% (14,396) of TE-related genes (Figure 3-A; Supplementary Table 3). In the KitaakeX genome, which has 35,594 annotated protein-coding genes (24), we identified 25,053 genes affected by mutations, representing about 70.38% coverage (Figure 3-A and Supplementary Table 3). Due to current limitations in KitaakeX genome annotation, a comparable detailed classification of affected genes into TE and non-TE categories was not feasible. This substantial gene coverage across both annotations underscores the extensive mutagenesis achieved in this study and highlights the resource’s power for genome-wide functional analysis.

**Figure 3.**
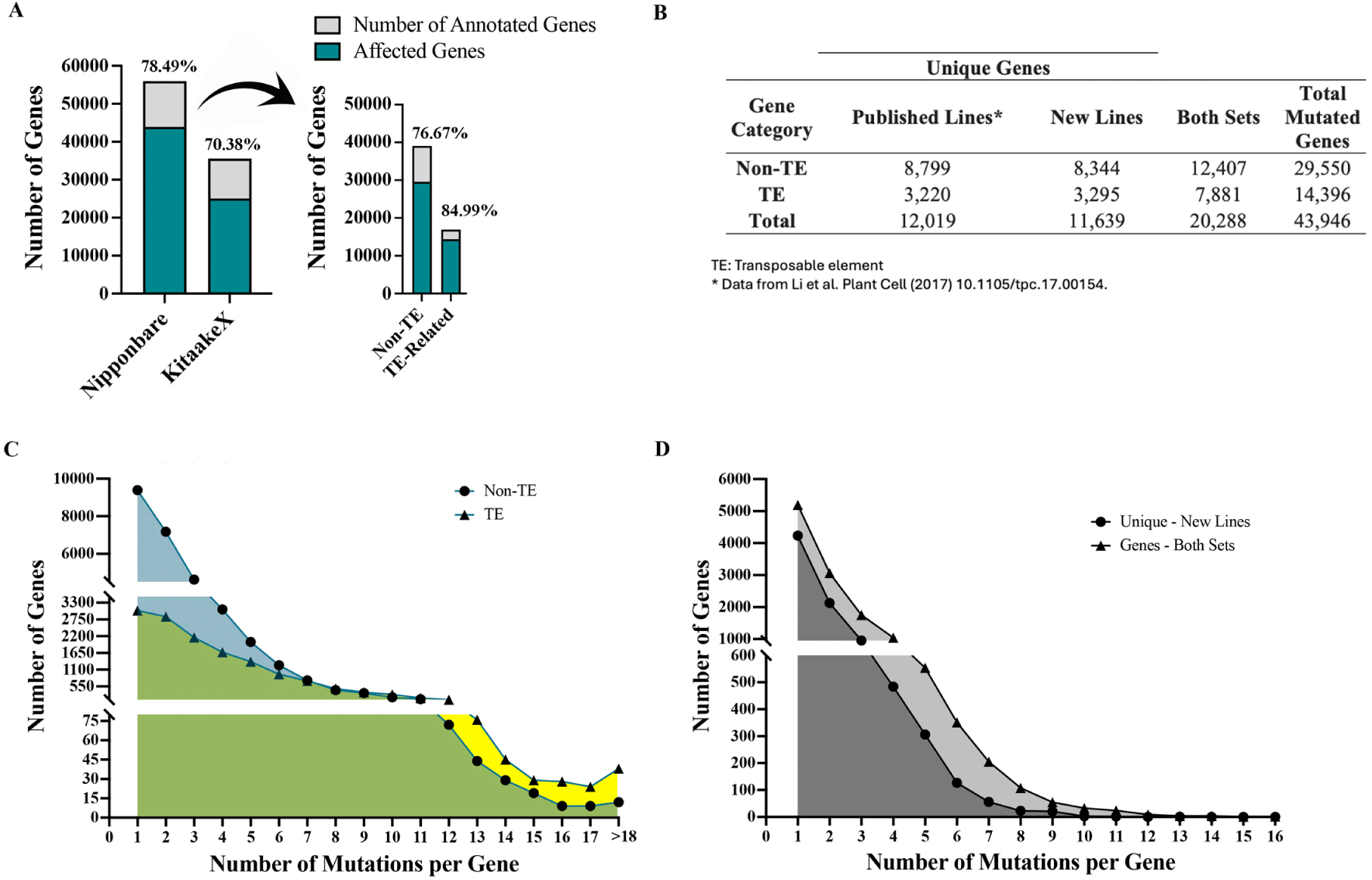
Comprehensive Analysis of Affected Genes in 3,268 FN-Induced Rice Mutant Lines. **(A)** Left: Proportion of annotated genes affected by fast-neutron-induced mutations in the Nipponbare and KitaakeX reference genomes, without subdivision of transposable element (TE) and non-TE gene categories. Right: Proportion of TE and non-TE genes affected within the Nipponbare reference genome. **(B)** Summary table of affected genes identified in each sequenced set based on the Nipponbare reference genome, including the number of genes unique to each set and those shared between both sets. Genes are further classified as TE or non-TE. **(C-D)** Frequency distribution of the number of mutations per gene in the Nipponbare reference genome. **(C)** Distribution across the entire KitBase population, with TE genes shown in yellow and non-TE genes in blue. **(D)** Distribution within the newly sequenced lines, with genes mutated in both the previous and new sets shown in dark grey and genes uniquely affected in the new set shown in light grey.

The expanded set of 1,764 new mutants significantly increased both gene coverage and allelic diversity beyond the original 1,504 lines. Among the Nipponbare-annotated genes, we identified 11,639 previously unaffected genes from the initial population (Figure 3-B and Supplementary Table 3). This increase comprises 8,344 (39.35%) non-TE genes and 3,295 (29.68%) TE-related genes. In addition, a total of 20,288 affected genes (12,407 non-TE and 7,881 TE) were identified in both sets. Analysis of mutation frequency across the entire KitBase population revealed that 68.22% of non-TE and 78.86% TE genes carry more than two independent mutations, with average mutation rates of 2.9 and 3.9 per gene, respectively (Figure 3-C and Supplementary Table 3). Within the newly sequenced lines alone, 49.24% of uniquely mutated non-TE genes harbor more than one mutation, with individual genes carrying up to 13 mutations. When considering genes mutated in both the previous and new sets, 58.14% of shared non-TE genes carry more than one mutation across the combined population, with some genes accumulating 16 independent mutational events (Figure 3-D and Supplementary Table 3). This variability indicates that individual genes may now harbor multiple distinct mutations across the entire population, potentially leading to diverse functional consequences and providing valuable allelic series.

To relate findings from the KitaakeX alignment to the better-annotated Nipponbare reference, we used two independent ortholog inference approaches (Best Hit in Rice and Inparanoid) to identify putative orthologous relationships among mutated genes. Of the 25,053 mutated genes identified in the KitaakeX alignment, 19,101 (76.24%) were concordantly assigned by both methods, 3,430 (13.69%) yielded discordant results, and 2,522 (10.07%) were not recovered by either analysis (Supplementary Table 3). Within the concordant set, 73 Nipponbare gene IDs were shared across multiple KitaakeX genes, yielding a final set of 19,028 unique Nipponbare gene IDs. Of these, 18,610 are non-TE genes, of which 16,791 (90.23%) were also found to be mutated in the Nipponbare alignment (Supplementary Figure 1-B and Supplementary Table 3). Comparative analysis of the full putative ortholog set revealed a high degree of overlap: 17,174 genes (90.26% of total mapped orthologs) were independently identified as mutated in the Nipponbare alignment (Supplementary Figure 1-C and Supplementary Table 3). This strong concordance between the two reference-based analyses, despite differences in absolute mutation counts, supports the reliability of the identified gene sets and increases confidence in the biological relevance of the reported mutations. It should be noted that this analysis summarizes putative ortholog relationships between Nipponbare and KitaakeX gene models, with emphasis on the top-ranked one-to-one ortholog assignments. For genes with ambiguous mappings, downstream analyses should also consider all additional ortholog candidates.

To further elucidate the chromosomal distribution and assess mutation coverage density within the affected genes, we mapped all identified mutations onto their respective chromosomal locations and determined the proportion of mutated genes per chromosome for both Nipponbare and KitaakeX genomes (Figure 4-A and Supplementary Figure 1-C). The coverage of mutated genes is very similar across chromosomes, with an average of 79.37% of genes mutated per chromosome in Nipponbare and 71.38% in KitaakeX (Table 1). For Nipponbare, the percentage of genes mutated on individual chromosomes ranged from 71.55% (chromosome 2) to 86.87% (chromosome 10), with most chromosomes showing saturation levels between 78% and 82%. The KitaakeX-based analysis revealed a similar pattern of coverage distribution across chromosomes, despite differences in absolute gene numbers due to annotation. Importantly, this analysis shows that no chromosome is left largely unmutated, indicating comprehensive coverage across the entire genome (Figure 4-A and Supplementary Table 3).

**Figure 4.**
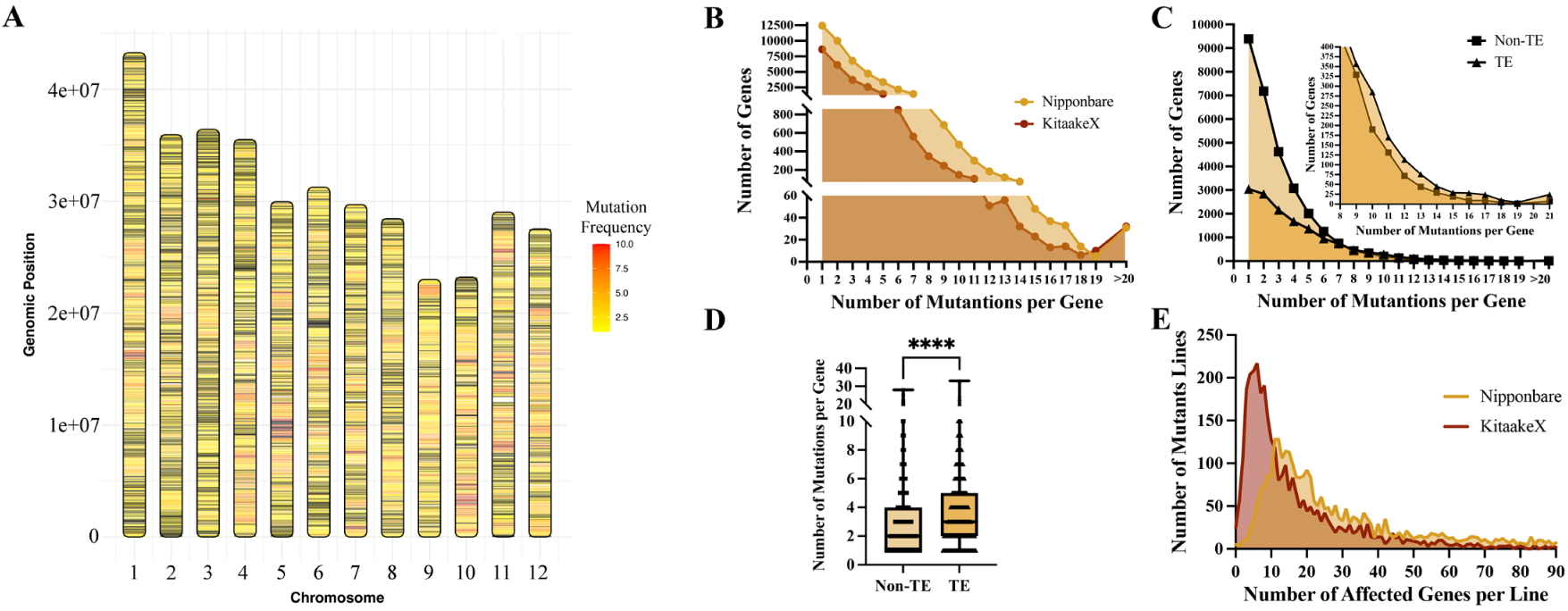
Chromosomal Distribution and Mutation Density of Affected Genes. **(A)** Chromosomal mapping of affected genes in the Nipponbare alignment. The heatmap represents the number of distinct mutations per gene, ranging from yellow (one mutation) to red (ten or more mutations). Non-mutated genes are shown in black, and intergenic regions are depicted in white. **(B)** Frequency distribution of mutations per gene across all affected genes in the Nipponbare and KitaakeX alignments, illustrating differences in mutation accumulation patterns between the two reference genomes. **(C)** Frequency distribution of mutation occurrences per gene in the Nipponbare alignment, stratified by gene classification (TE and non-TE genes). **(D)** Boxplot of the number of mutations per gene in the Nipponbare alignment, stratified by gene classification (TE and non-TE genes). Statistical significance was assessed using a two-sample t-test. **(E)** Distribution of the number of affected genes per mutant line for both the Nipponbare and KitaakeX alignments. Data shown represent more than 90% of all affected genes per line; complete data are provided in Supplementary Table 1.

To characterize the frequency of mutations within affected genes, we analyzed the distribution of mutation occurrences per gene across all affected genes in both reference alignments. In the Nipponbare alignment, 12,435 of 43,946 affected genes (28.3%) carried a single mutation, while the majority, 31,511 genes (71.7%), harbored two or more independent mutations (Figure 4-B and Supplementary Table 3). When stratified by gene classification, non-TE genes showed a higher proportion of genes with low mutation counts compared to TE genes, with TE genes exhibiting a progressive enrichment toward higher mutation frequencies (Figure 4-C). This difference in mutational burden between TE and non-TE genes was statistically significant (Figure 4-D), suggesting that, despite the random nature of fast-neutron irradiation, TE-associated regions accumulate mutations at a higher rate than non-TE genes. A similar trend was observed in the KitaakeX alignment, where 8,616 of 25,053 affected genes (34.4%) carried a single mutation and 16,437 (65.6%) harbored two or more mutations. This multiplicity of independent mutational events within individual genes constitutes a valuable resource, enabling cross-validation of gene function across multiple alleles and providing the means to substantiate or refute the association between a given gene and a specific phenotype observed across different mutant lines.

The average number of affected genes per line was 42.52 when aligned to the Nipponbare reference genome and 21.33 with the KitaakeX reference (Figure 4-E and Supplementary Table 1). This discrepancy could be primarily correlated with the differences in gene annotations between the two genomes; the more comprehensively annotated Nipponbare genome encompasses a larger set of features than KitaakeX, resulting in more mutations falling within annotated gene boundaries. The distribution analysis revealed that 90% of the lines have fewer than 88 affected genes in the Nipponbare alignment and fewer than 35 in the KitaakeX alignment (Figure 4-E and Supplementary Table 1). Focusing on this subset, the average number of disrupted genes per line decreases to 27.09 for Nipponbare and 12.2 for KitaakeX. These numbers underscore the utility of this mutant population for functional genomics and facilitating genetic segregation studies. Notably, 2,875 lines exhibited a higher number of affected genes per line in the Nipponbare alignment compared to KitaakeX, while 49 lines showed identical counts in both alignments (Figure 4-E and Supplementary Table 1).

### Transcription Factor Coverage

Transcription factors (TFs) are key regulatory genes that control fundamental plant processes. Mutant collections with extensive coverage of TF families are particularly useful for understanding regulatory networks (40,41). We examined how many of the annotated rice TF genes are disrupted in the KitBase population. Based on the Nipponbare genome annotation, which contains 1,862 putative TF genes (35), we found that 1,385 TF genes carry mutations in our population, representing substantial coverage of 74.4% of all TFs (Supplementary Figure 1-D; Supplementary Table 3). This indicates that FN mutagenesis broadly impacted regulatory genes, and most TF families have multiple members mutated. Indeed, all major TF families (AP2/ERF, bHLH, MYB, bZIP, NAC, etc.) have at least 50% of their genes mutated (Supplementary Figure 1-E; Supplementary Table 3). We observed particularly good coverage for many families, with, for example, over 70% of homeobox and MADS-box genes mutated in KitBase. The sole exception was the small S1Fa-like family, where none of the few members had a mutation. This extensive collection of TF mutants provides a powerful means to study gene regulatory networks controlling crucial aspects of plant development, metabolism, and stress responses.

### Characterization of Non-Mutated Genes in KitBase

To comprehensively understand the full impact of FN-irradiation on the rice genome and leverage the extensive coverage achieved in our population, we analyzed genes that remained unmutated across all 3,268 lines in the KitBase resource. Genes that remain unmutated in a large, randomly mutagenized population like this are strong candidates for essential genes, often involved in critical biological processes. Disruption of such genes frequently leads to gametophytic or sporophytic lethality, or severe developmental defects that prevent the recovery of viable homozygous mutant plants in the second (M2) generation, thus explaining their absence from a screen of viable M2 lines (42).

We identified 11,855 genes in the Nipponbare reference for which no mutation was detected in any of the KitBase lines (Supplementary Table 3). Among these, 9,528 are non-TE genes, and 2,327 are TE genes. A similar analysis in KitaakeX yielded a list of 9,585 non-mutated genes; this smaller number compared to Nipponbare likely reflects the annotated gene set. While the absence of mutations in some genes, particularly those with small coding regions, could theoretically be due to chance, in a large and highly saturated population like KitBase, the persistent lack of detected mutations in a gene strongly indicates its potential essential nature. Knockouts of essential genes often lead to severe deleterious effects, which prevent the recovery of viable mutant progeny (42). Similar patterns have been observed in other organisms, where essential genes are under-represented in mutant collections because null mutations cannot be propagated (43).

To further investigate the nature of these non-mutated genes and explore the essential gene hypothesis, we analyzed their expression patterns and functional categories using data from the Nipponbare alignment. An *in silico* expression survey using RNA-seq data from 13 rice tissues revealed that the non-mutated genes tend to be expressed (Supplementary Figure 1-F and Supplementary Table 3). Specifically, over 33% of the non-mutated non-TE genes showed moderate expression (10–50 FPKM) in at least one tissue, and 575 genes (6.72%) exhibited high expression (average >50 FPKM). Surprisingly, among the highly expressed subset, 61 genes were annotated as ’expressed protein’, indicating they are actively transcribed but their specific function remains unknown. The presence of highly expressed genes may indicate possible essential functions, where mutations could be deleterious to plant survival, thereby being negatively selected during mutagenesis.

Functional enrichment analysis of the highly expressed non-mutated genes showed a significant over-representation of genes involved in fundamental biological processes. These include translation, primary metabolic processes (e.g., ribosomal proteins, core enzymes), biosynthetic processes, and protein metabolic processes (Supplementary Figure 1-G and Supplementary Table 3). Complementary pathway enrichment analysis further highlighted the participation of these genes in key biological pathways critical for survival, including ribosome function, core metabolic pathways, oxidative phosphorylation, and protein processing in the endoplasmic reticulum (Supplementary Figure 1-H and Supplementary Table 3). Collectively, these results indicated that this subset of highly expressed genes may be essential for plant survival and thus resistant to knockout mutagenesis. This implies that loss-of-function in those genes is likely lethal or strongly selected against in the developing plant, preventing their representation in the viable M2 population.

While a subset of unmutated genes are strong candidates for essentiality, it is important to note that some genes classified as non-mutated here could potentially be mutable if we screened an even larger population. Approximately 59.6% of these non-mutated genes showed low or background-level expression (FPKM <10), suggesting they may have been missed by chance due to their small coding regions or low mutation probability. Supporting this, our analysis showed that non-mutated genes with low or background expression are, on average, 50.52% shorter in nucleotide length compared to genes with moderate to high expression. Likewise, genes with no expression data were 32.65% shorter (Supplementary Figure 1-I; Supplementary Table 3). This suggests that increasing the population size could potentially lead to the identification of mutations in this subset of genes.

Genomic mapping demonstrated a uniform distribution of non-mutated genes across all chromosomes, with no significant clustering (Supplementary Figure 1-J). Focusing on the highly expressed unmutated genes (likely essential candidates), we observed variations in their counts per chromosome. Chromosomes 2, 3, and 1 harbored the highest numbers (113, 108, and 80 genes, respectively), while chromosomes 9 and 12 contained the fewest (2 and 17 genes) (Supplementary Figure 1-I and Supplementary Table 3). Additionally, examination of the genomic landscape revealed interesting features within the unmutated gene set, such as the presence of very short intergenic distances between some unmutated genes on chromosomes 10 and 3 (26 bp and 71 bp, respectively). These findings offer valuable insights into the resilience of certain genes to mutagenesis. Importantly, our results present a list of likely essential genes in rice, which can be useful for future functional genomics studies.

### Mutation Spectrum and Types of Variants

Fast-neutron mutagenesis is known to induce a broad spectrum of genetic alterations, including deletions, insertions, inversions, translocations, single-base substitutions (SBS), and tandem duplications (23). Identifying the types of mutations is crucial for predicting their impact on gene function. In this study, we characterized the types and distributions of mutations identified through alignments with Nipponbare and KitaakeX reference genomes (Supplementary Table 2).

Upon mapping the different types of mutation across the chromosomes, we observed a uniform distribution, indicating the absence of mutational hotspots (Figure 5-A). In the Nipponbare-aligned, SBS was the most abundant class, constituting 106,232 mutations (46.07% of the total) (Figure 5-B and Supplementary Table 2). Deletions were the next most common, with 75,808 events (32.88%), followed by insertions (20,461 mutations, 8.87%). We also identified 16,119 putative translocations (6.99%), 11,859 inversions (5.14%), and 65 tandem duplications (0.0028%). The KitaakeX-aligned showed a similar distribution: SBS accounted for 116,875 mutations (55.1%), deletions 59,619 (28.1%), insertions 12,196 (5.75%), inversions 16,208 (7.64%), and translocations 7,183 (3.38%) (Figure 5-A and B; Supplementary Table 2). Notably, tandem duplications were only identified in the original analysis of the 1,504 Nipponbare-alignment lines and were not analyzed in the newly sequenced lines or within the KitaakeX pipeline. Despite these nuances in variant calling, the combined analysis across both references contributes to a more complete catalog of the diverse mutation spectrum induced by fast neutrons.

**Figure 5.**
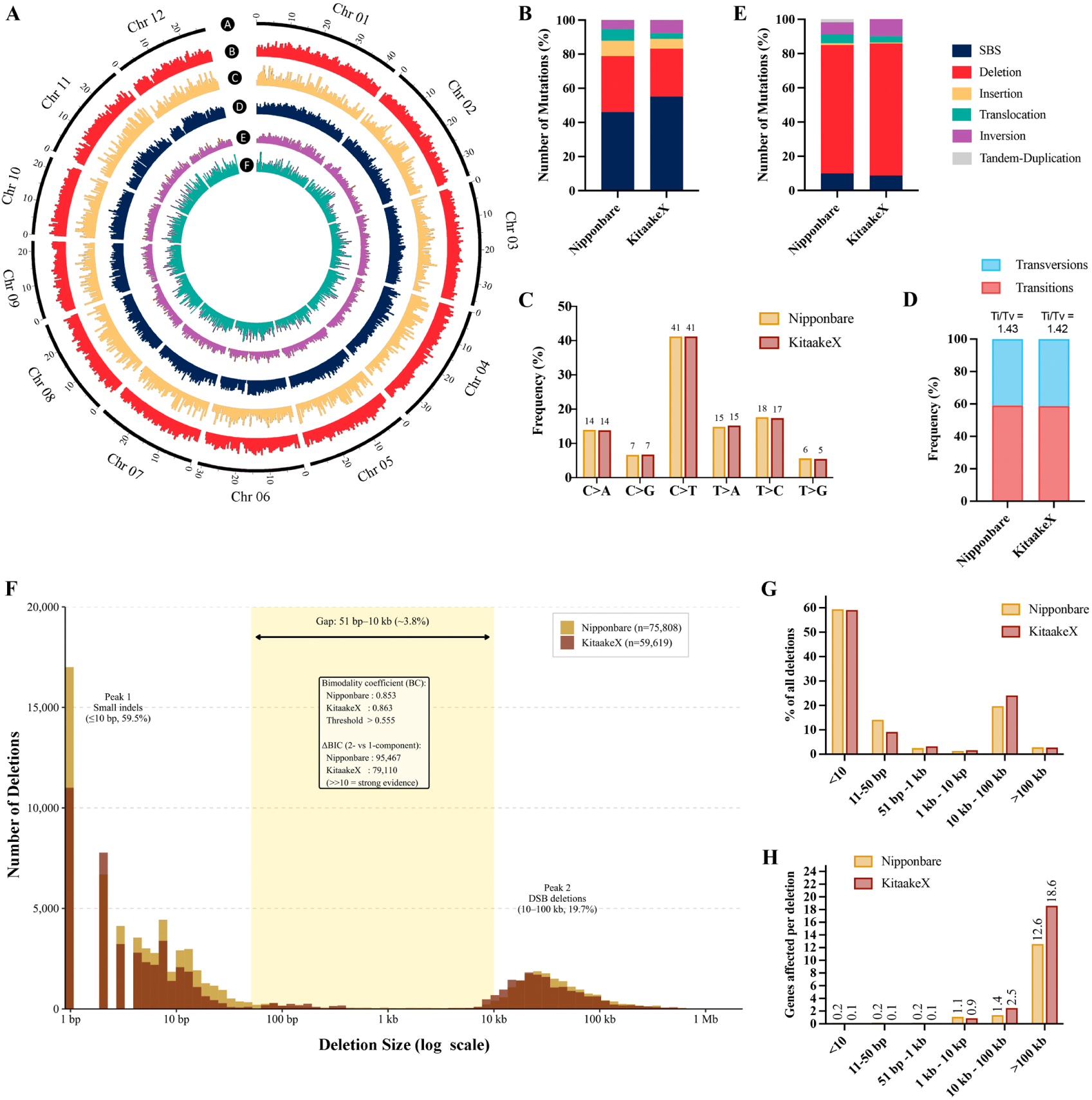
Chromosomal Distribution and Frequency of Mutation Types in the FN-Induced Kitaake Rice Mutant Population. **(A)** This figure illustrates the distribution and frequency of various mutation types across the ‘A’ 12 rice chromosomes, presented on a megabase scale. Each panel highlights a specific mutation type with color-coded representations to enhance clarity. ‘B**’** Deletions are depicted in dark red, indicating regions where nucleotide sequences have been removed. ‘C’ Insertions are shown in yellow, marking locations where additional nucleotide sequences have been incorporated. ‘D’ Single-base substitutions (SBS) are represented in dark blue, denoting points where a single nucleotide has been altered. ‘E’ Inversions are illustrated in purple, highlighting segments where the orientation of a DNA fragment has been reversed. ‘F’ Translocations are presented in marine green, identifying instances where DNA fragments have been relocated to different positions within the genome. **(B)** Bar chart depicting the overall frequency of each mutation type across the entire genome in both alignments. **(C)** Distribution of SBS frequencies using the standard pyrimidine-reference SBS6 classification for mutations aligned to the Nipponbare and KitaakeX reference genomes. The six canonical substitution classes (C>A, C>G, C>T, T>A, T>C, T>G) each represent complementary strand pairs: for example, C>T includes both C>T and G>A events. Solid bars represent Nipponbare alignment; hatched bars represent KitaakeX alignment. **(D)** Proportion of transitions (Ti) and transversions (Tv) in each alignment, with Ti/Tv ratios indicated above each bar. **(E)** Bar chart illustrating the frequency of each mutation type specifically within affected genes. **(F)** Log₁₀-scale histogram of deletion sizes for all deletions identified in the Nipponbare (dark gold) and KitaakeX (dark red) reference alignments. The yellow shaded region indicates the gap zone (51 bp – 10 kb). **(G)** Percentage of total deletions in each size class for the Nipponbare (dark gold) and KitaakeX (dark red) alignments. Six size classes are shown: ≤10 bp, 11–50 bp, 51 bp–1 kb, 1–10 kb, 10–100 kb, and >100 kb. **(H)** Average number of genes affected per deletion event for each size class, shown for the Nipponbare (dark gold) and KitaakeX (dark red) alignments. Gene disruption rates were calculated by intersecting deletion coordinates with gene body annotations from Supplementary Table 3.

Overall, a key observation is that SBS and deletions collectively represent the majority of FN-induced mutations, accounting for approximately 80% of all detected variants (Figure 5-B and Supplementary Table 3). Upon detailed analysis of SBS, we observed that both alignments exhibit a similar mutation distribution pattern, revealing a highly consistent substitution spectrum across both reference genomes (Figure 5C-D). Using the standard pyrimidine-reference SBS6 classification, C>T transitions predominated, accounting for 41.5% of all SBS in both alignments, more than twice the frequency of any other substitution class. T>C transitions were the second most frequent (17.5% and 17.2%), followed by C>A transversions (14.1% and 14.0%). Overall, transitions accounted for 59.0% and 58.7% of SBS respectively, yielding transition-to-transversion (Ti/Tv) ratios of 1.44 and 1.42, significantly above the Ti/Tv of ∼0.5 expected under random mutation. The near-identical spectra between the two reference alignments confirm that this mutational signature is a robust property of fast-neutron irradiation, independent of reference genome choice. At the gene level, TE-associated genes showed a significantly elevated C>T frequency compared to non-TE genes (47.9% vs. 37.3%, χ² = 101.7, p = 2.3×10⁻²⁰), consistent with hypermutation at methylated cytosines characteristic of transposable element regions (Supplementary Figure 1-L/M).

Another major class of mutations prevalent in the FN population is deletions. To characterize their size distribution and impact, we analyzed all identified deletion events (Table 2). In both the Nipponbare and KitaakeX alignments, the vast majority of deletions were relatively small (under 100 bp), representing 74.8% and 69.4% of all deletions, respectively. Single-base deletions alone comprised a significant proportion, accounting for 22.4% of all deletions in Nipponbare and 18.5% in KitaakeX. At the other end of the spectrum, FN mutagenesis also produces a number of large deletions; we observed many deletions in the 1 kb–100 kb range, and a few in the megabase range. The largest deletion spanned approximately 18 Mb in one line, essentially removing a large chromosome segment, as previously noted (16). When deletion sizes were visualized on a log₁₀ scale to resolve the full six-order-of-magnitude range, a striking bimodal distribution emerged that is completely obscured on a linear scale (Figure 5-F). A prominent first peak centered on micro-indels (≤50 bp, 73.6% of all deletions) was separated from a second peak of large deletions (10–100 kb, 19.7%) by a markedly underrepresented gap spanning 51 bp to 10 kb, which contained only 3.8% of Nipponbare deletions and 3.2% of KitaakeX deletions (Figure 5-G). This bimodal structure was formally confirmed by bimodality coefficient analysis (BC = 0.853 and 0.863 for the two alignments; threshold > 0.555) and Gaussian mixture modeling (ΔBIC > 79,000 in both alignments, far exceeding the threshold of 10 for strong evidence of two components). The consistency of this pattern across both reference genomes confirms it is a biological property of FN mutagenesis and not na alignment artifact. Mechanistically, small deletions (≤50 bp) are consistent with non-homologous end joining (NHEJ) microresection and replication slippage, while the large deletion peak (10 -100 kb) reflects double-strand breaks (DSBs) repaired by microhomology-mediated end joining (MMEJ) or complete repair failure, the primary physical mechanism of fast-neutron DNA damage (16,63). The pronounced gap between these two peaks suggests these mechanisms operate largely independently, a pattern consistent with fast-neutron-mutagenized Arabidopsis thaliana (63). While the average deletion size was 13.6 kb (Nipponbare) and 14.5 kb (KitaakeX), both means are heavily skewed by the few extremely large events. The median deletion size was 7 bp in both alignments, underscoring that the vast majority of FN-induced deletions show to be small, or in another way, precise events.

When focusing on mutation events occurring within gene regions, deletions emerged as the most prevalent mutation type affecting gene sequences (Figure 5-D and Supplementary Table 3). In the Nipponbare alignment, we identified 19,591 deletion events, mutating 38,961 unique genes (88.7% of all affected genes). In the KitaakeX alignment, 13,616 deletions affecting 21,630 (86.3% of the affected genes). Despite SBSs being the most abundant mutation type overall, they affected a smaller proportion of genes, 9,933 genes (22,6%) in Nipponbare and 4,881 genes (19.5%) in KitaakeX. This discrepancy suggests that deletions often have a more profound impact on gene integrity due to their potential to remove entire gene sequences or regulatory regions. The two deletion size classes identified above differ dramatically in their gene disruption impact (Figure 5-H): small deletions (≤10 bp) affect na average of only 0.15 genes per deletion event in the Nipponbare alignment (0.10 in KitaakeX), providing surgical, largely single-gene knockouts ideal for unambiguous genotype-phenotype analysis. By contrast, deletions in the 10 - 100 kb class affect 1.36 genes per event on average (2.49 in KitaakeX), and those exceeding 100 kb affect an average of 12.6 genes per event (18.6 in KitaakeX), enabling the study of gene redundancy and contiguous gene cluster function by disrupting multiple adjacent genes in a single event (44,45). Both deletions and point mutations induced by various mutagens can increase the likelihood of uncovering desirable traits that were previously suppressed during selective breeding due to linkage drag (9).

### Phenotypic Diversity in the Mutant Population

Phenotypic characterization of sequenced mutant lines is a valuable resource for rice genetics, enabling researchers to efficiently associate observable traits with underlying genetic mutations. This integrated genomic and phenotypic data specifically facilitates forward genetic analyses, allowing for the identification of genes responsible for specific phenotypic variations based on sequence data. To support such analyses, a set of individual mutagenized lines was systematically phenotyped for a range of agronomically relevant traits, expanding the functional utility of KitBase.

We conducted systematic phenotypic characterization on a large subset of over 2,700 mutant lines (M₂ or M₃ generation) grown under normal conditions. This effort focused on a set of readily quantifiable and agronomically relevant traits, defined using standard ontology terms, including germination rate (seedling vigor), albino seedling frequency, tiller number, days to heading (flowering time), plant height, panicle traits (panicle length/weight and filled grain number), and seed traits (seed number per panicle, seed fertility) (Figure 6 and Supplementary Table 4). Phenotypic data for each line were recorded for one or more of these traits and compared to wild-type Kitaake grown in the same environment. To facilitate comparison across different planting batches and minimize the impact of environmental variation, trait values were normalized relative to the wild-type control (KitaakeX), where a value of 1 or percentage representation (100%) represents wild-type performance (Supplementary Figure 2-A and Supplementary Table 4). This normalization approach ensures trait data from diverse seasons or greenhouse conditions are comparable on a consistent scale, enhancing the data’s utility for genetic analysis.

**Figure 6.**
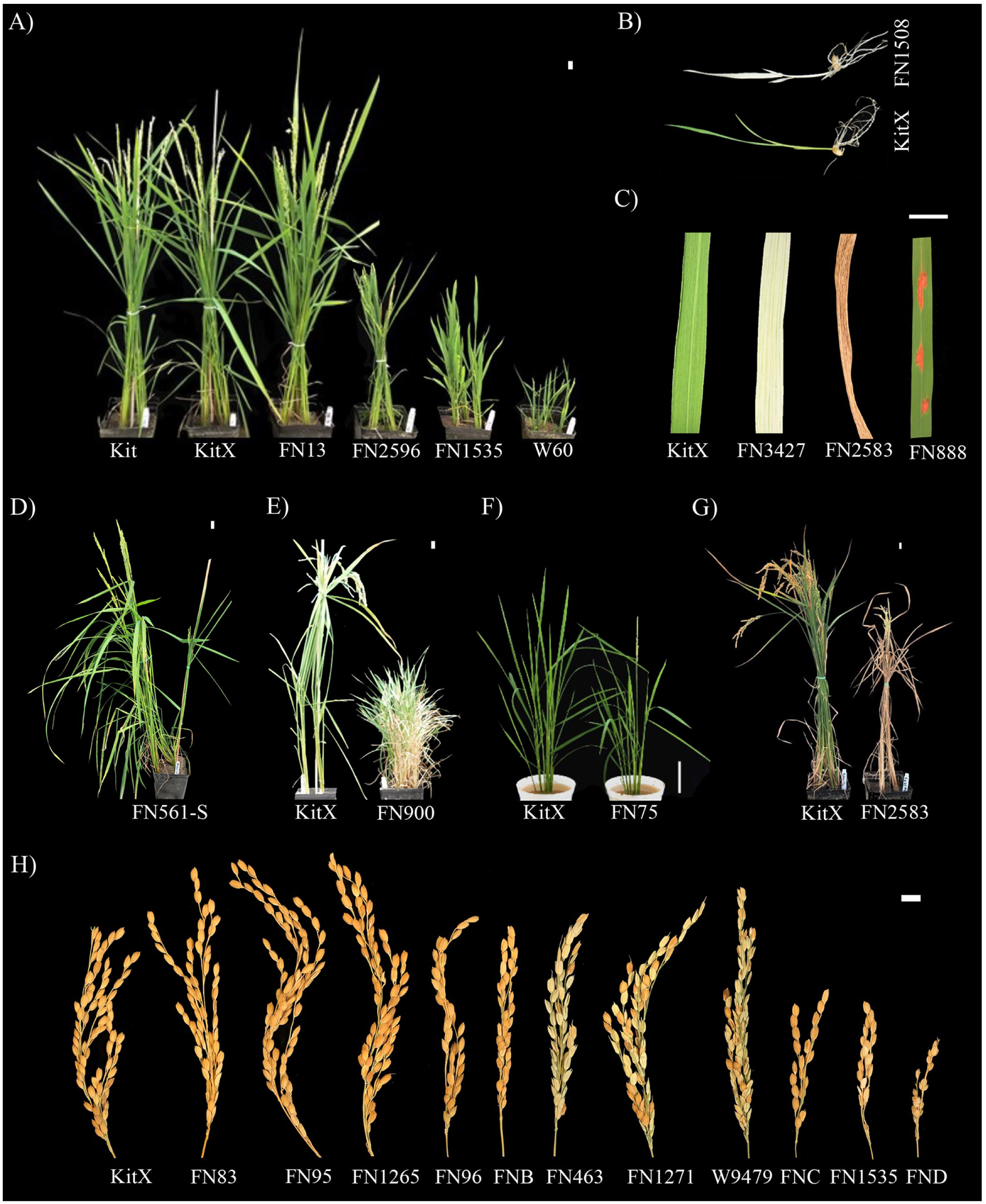
Representative phenotypes observed in fast-neutron-mutagenized KitaakeX rice lines. This figure displays a range of morphological alterations induced by FN mutagenesis in the KitaakeX population. A) Panicle height: tall to dwarf; B) Lethal Albino seedling; C) Leaf Color: partial albino, brown and mimic-lesion; Number of Tillers: D) Reduced and E) Enhanced; F) Days to Heading: Early flowering; G) Early senescence; H) Different panicle morphology: panicle length, panicle fertility, number of seeds per panicle. These images highlight the extensive phenotypic diversity generated by fast-neutron irradiation. KitX = KitaakeX; FNB = FN1135; FNC = FN1299; and FND = FN1727.

Phenotypic analysis revealed substantial variation among the mutant lines across all evaluated traits, demonstrating the power of FN mutagenesis to generate diverse functional alterations (Figure 6, Supplementary Figure 2-A, and Supplementary Table 4). For example, days to heading ranged from 21% earlier (average of 41 days in FN1396-S) to over 34% later (average of 71 days in FN2092-S) compared to the KitaakeX control. Plant height varied widely, from a 71.5% reduction (26.7 cm in w60-2-13) to an 8.1% increase (101.6 cm in FN1016-S). Four lines were categorized as short, averaging 54% shorter than KitaakeX, while 58 additional lines exhibited a dwarf phenotype without height quantification. Mutants affecting seedling development were also observed, with 33 lines showing segregation for an albino seedling phenotype among the progeny of sequenced, phenotypically WT (green) individuals. This suggests that the sequenced plants likely carried the causal mutation in a heterozygous state. Tillering capacity showed a broad range: 54 lines displayed a low tiller number (average of 3), contrasted by another 54 lines with high tillering (average of 6 tillers). Regarding seed yield, four lines were completely sterile (FN1015-S, FN1022-S, FN4068-S, and FN4445-S), whereas 116 lines were classified as highly productive, with yields averaging 120% greater than KitaakeX, suggesting the presence of potentially beneficial mutations in the population.

To elucidate the relationships among various phenotypic traits evaluated, we conducted a correlation analysis. This analysis revealed 15 traits with statistically significant associations. Among these, four traits (panicle weight, plant height, seed number, and number of empty seeds) exhibited moderate to strong correlations (Supplementary Figure 2-B; Supplementary Table 4). Subsequently, principal component analysis (PCA) was performed, which effectively differentiated subsets of lines based on their overall phenotypic profiles. PCA particularly highlighted variations related to seed yield and plant height as major components distinguishing lines (Supplementary Figure 2-C). These findings on phenotypic relationships and major sources of variation align with prior research demonstrating the utility of correlation analysis and PCA in distinguishing rice lines based on key agronomic traits. Such analyses help users of the KitBase resource understand the structure of the phenotypic data and can inform gene discovery efforts by identifying potentially linked traits or major phenotypic classes.

In addition to the quantitative traits evaluated, we also cataloged various qualitative phenotypic variations observed within the mutant population (Supplementary Table 4). These include alterations in leaf color (e.g., lighter green, yellow, white stripes), leaf morphology (e.g., curling and lesion mimic), panicle architecture (e.g., variations in grain size, presence of long awns, stunted growth), and developmental anomalies (e.g., robust stems, brown spots, brown roots, early senescence, sterility in segregation, reduced tiller number). To further aid resource users, we captured representative photographs of some mutants exhibiting specific traits and have made these images available on the KitBase website, linked to the corresponding genomic data. The sheer diversity of these qualitative phenotypes, complementing the quantitative data, further underscores the extensive functional genetic variability induced by fast-neutron mutagenesis within the KitBase population.

### KitBase Web Interface and Data Access

Public access to well-structured, high-throughput genomic and phenotypic resources is essential for advancing scientific research and accelerating discoveries. To facilitate public use of the KitBase resource and its integrated datasets, we have updated the KitBase web interface (http://kitbase.ucdavis.edu/) with new features for data query and visualization (Figure 7-A). Users can easily search the database for relevant mutant lines by various criteria, including Line ID, Gene ID, Keyword, or specific Phenotypic trait (Figure 7-B). The results of these searches provide integrated genomic and phenotypic information, presented in user-friendly, detailed tables and pages (Supplementary Figure 3, A-D). These updates provide the research community with efficient and intuitive methods to locate and explore lines, genes, or traits of interest within the KitBase population.

**Figure 7.**
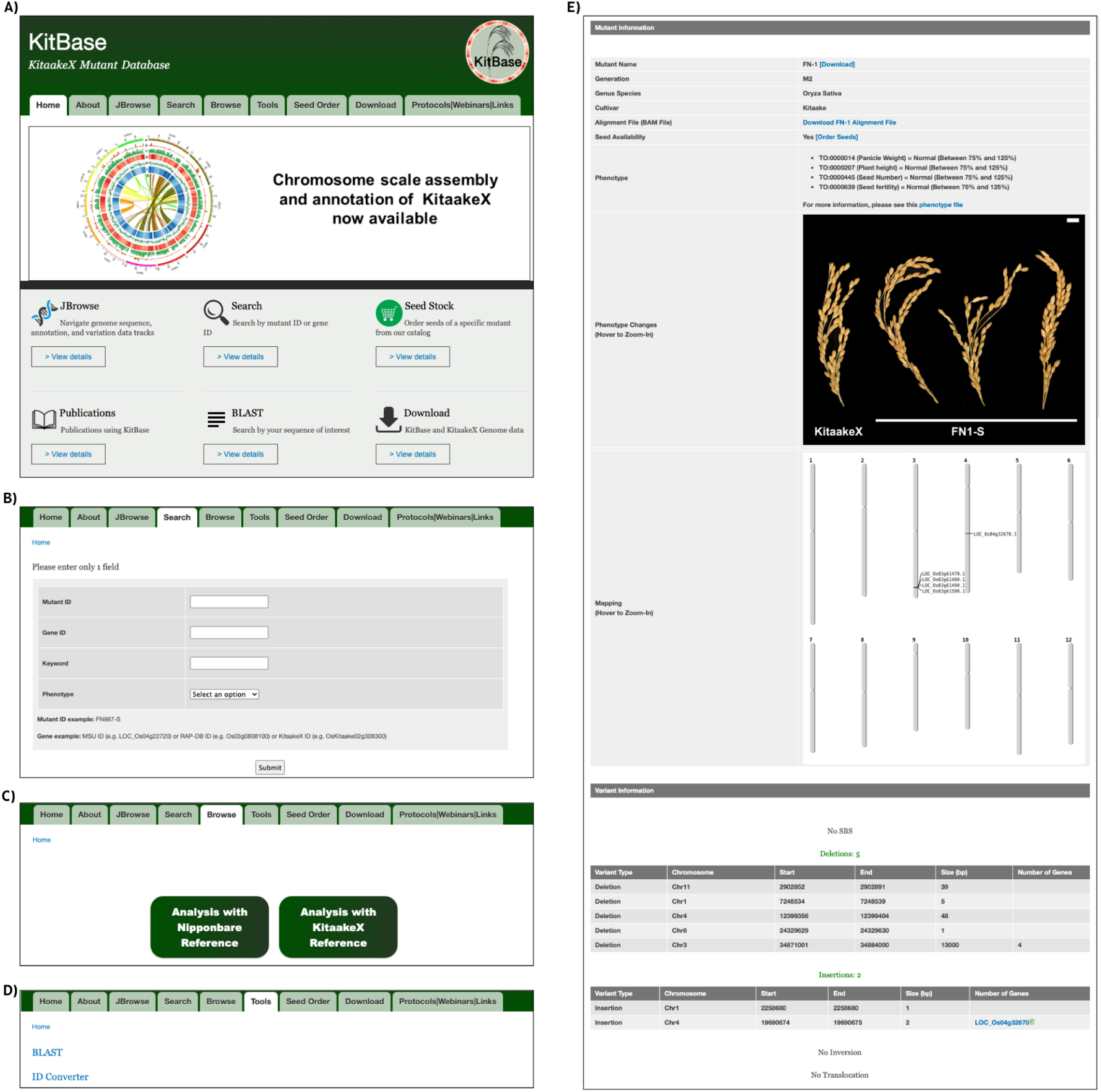
Overview of the KitBase User Interface. **(A)** Main Navigation Page: Displays the primary navigation menu at the top and bottom, facilitating access to various sections of KitBase. **(B)** Search Functionality: Offers multiple search options, including Mutant ID, Gene ID, Keyword, and Phenotype, enabling users to efficiently locate specific data within the database. (**C)** Browse Interface: presents two distinct alignment options. Selecting an alignment reveals detailed data for the corresponding mutant lines. **(D)** Tools Section: Features utilities such as BLAST and an ID converter between Nipponbare and Kitaake gene IDs, and vice versa. **(E)** Mutant Line Information: Provides comprehensive details on individual mutant lines, including genotype and phenotype data.

To provide comprehensive access and utility, all the sequencing data generated for both Nipponbare and KitaakeX alignments are freely accessible for download via the online platform (Figure 7-C). This enables researchers to perform their own custom analyses. Furthermore, the ’Tools’ section of the website includes valuable bioinformatics functionalities (Figure 7-D and Supplementary Figure 3-F). These tools include BLAST, allowing users to search for sequence similarities within the KitBase data, and an ID converter, which is particularly useful for identifying possible orthologous genes between Nipponbare and Kitaake gene IDs and bridging annotation differences (Supplementary Figure 3-G).

Each mutant line in KitBase is provided with a dedicated page containing comprehensive, integrated information (Figure 7-E). This page includes information about the sequence, phenotype, and genotype data. In addition, each page provides access to sequenced data, allowing users to download and examine predicted mutations. Furthermore, the new JBrowse genome browser is integrated into the platform, allowing for interactive visualization of mutations in the KitBase population in the Nipponbare reference-genome context, aligned along each chromosome (Supplementary Figure 3-E). JBrowse can be particularly useful for identifying multiple lines with mutations in the same gene or overlapping genomic region. It also facilitates the examination of mutations located in intergenic or regulatory regions that may not be immediately linked to an annotated gene, supporting the identification of regulatory mutations.

For researchers interested in obtaining specific mutant lines, the KitBase website includes a dedicated ‘Seed Order’ page that provides a straightforward form to request the desired lines. To further support the research community, the website also offers resources in the ‘Download’ and ‘Protocols/Webinars/Links’ sections, with detailed protocols and instructional videos providing guidance on how to utilize the KitBase resource and its online tools. Collectively, the design and feature improvements of the KitBase web interface are specifically aimed at maximizing the accessibility and utility of the entire KitBase population.

### Impact of KitBase in the scientific society: Forward and reverse analysis

The KitBase resource has already proven invaluable for advancing functional genomic research in rice, facilitating the identification of genes controlling key traits. For example, a screen of a subset of the KitBase population successfully identified a mutant in the histidine kinase-1 (HK1) gene, which exhibited defective root circumnutation (46). Additional studies have demonstrated the utility of KitBase for researching the genetic basis of complex traits. In a separate grain morphology study, the *gs9–1* mutant allele, harboring a 3-bp deletion in the gene LOC_Os09g02650 (an allele of *BC12/GDD1/MTD1*), was identified within the KitBase collection (47). This specific mutation in *gs9–1* was shown to result in altered grain shape, reduced cell number and length in grain glumes, and defects in gibberellin biosynthesis, consequently affecting overall plant stature and yield.

The KitBase resource has also proven to be valuable for advancing disease resistance studies in rice. Mutants with altered XA21-mediated immunity were identified in the population, such as the *sxi2* mutant, which was found to harbor a 20-kb deletion removing the *PALD* gene (48). Similarly, the *sxi4* mutant, also showing altered XA21-mediated immunity, was identified with a 32-kb translocation affecting the gene encoding Dicer-like protein 2a (*DCL2a*) (49). These examples demonstrate the utility of KitBase in uncovering the genetic basis of rice immunity and highlight the resource’s capacity to provide structural mutations impacting key defense components.

KitBase has also been effectively used by researchers to isolate lesion-mimic mutants (LMMs) exhibiting enhanced disease resistance (50). Leveraging the mutant population, it was possible to identify a specific mutant harboring a 29-bp deletion in the *RESISTANCE TO BLAST1* (*RBL1*) gene. This gene encodes a cytidine diphosphate diacylglycerol synthase involved in phospholipid biosynthesis. The identified mutation in *RBL1* was shown to cause the lesion-mimic phenotype and enhanced disease resistance, highlighting *RBL1* as a novel broad-spectrum disease-resistance gene in rice.

To illustrate how KitBase integrates phenotypic and genomic data for rapid gene-phenotype validation, we screened the population for dwarf mutants by compiling 158 genes reported to possibly alter plant height in rice and querying KitBase for lines with mutations in these genes (Supplementary Table 5). We identified 263 FN candidate lines. Among these, *D1/RGA1* (LOC_Os05g26890), encoding a Gα subunit involved in gibberellin signaling, may be disrupted in 8 independent lines, two of which already showed segregation for dwarfism in KitBase phenotypic records (Supplementary Table 4). Line FN1535-S, reported previously (23), carries a chromosome 5 inversion truncating *D1/RGA1* within exon 4. A KitBase search immediately identified FN3664-S, harboring an independent 136-kb deletion (Chr5: 15,481,001-15,617,000 bp) that removes the entire locus (Supplementary Figure 4-A; Supplementary Table 5). Critically, *D1/RGA1* was the only gene mutated in both lines. In the FN3664-S segregating population, dwarfs appeared at a ∼1:3 ratio, and PCR genotyping confirmed cosegregation of the deletion with the dwarf phenotype (Supplementary Figure 4-B). Both alleles produced statistically indistinguishable phenotypes: at Day 40, FN3664-S Dwarf and FN1535-S plants reached ∼57% of wild-type height and showed equivalent reductions in panicle and seed dimensions (Supplementary Figure 4C-G). Together, these two independent alleles, identified using KitBase, provide reciprocal genetic confirmation that *D1/RGA1* disruption causes dwarfism, demonstrating how the resource accelerates validation without map-based cloning (Supplementary Figure 4H-J).

Beyond traditional forward genetics, the KitBase resource has also facilitated the development of advanced research methodologies and fundamental biological insights. For instance, the phenotypic diversity generated by KitBase’s induced mutations has been incorporated into training datasets for deep learning-based SLEAP frameworks (51). This application enabled the automated detection and localization of key root landmarks, allowing for the extraction of quantitative traits critical for genotypic classification and phenotypic mapping, thereby significantly enhancing high-throughput root phenotyping capabilities. Furthermore, the comprehensive genomic data from KitBase has facilitated fundamental studies on mutation rates and DNA repair. Reanalysis of the high-resolution *de novo* single-base substitution data from the FN lines revealed that genomic regions enriched for the epigenetic mark H3K4me1 exhibited significantly lower mutation rates (52,53). These findings suggest a conserved, epigenome-targeted DNA repair mechanism modulating mutation rate variation in plant genomes. In summary, as demonstrated by the diverse examples, KitBase has been proven to be a powerful and versatile genomic resource.

## Discussion

In this study, we significantly expanded and comprehensively characterized the KitBase, a FN-induced mutant population in rice, creating a valuable resource for functional genomic research. The addition of 1,764 newly sequenced lines to the original 1,504 (23) results in a robust repository of 3,268 mutant lines, extensively characterized at both the genomic and phenotypic levels (Figure 2-A, Supplementary Table 1). This expansion, coupled with a dual-reference genome alignment strategy and an updated, user-friendly web interface, collectively enhances the utility and accessibility of the KitBase platform for the global rice research community.

The expanded KitBase population offers exceptional genome coverage. Alignment to the well-annotated Nipponbare genome shows that approximately 78% of all annotated genes carry at least one mutation (Figure 3-A, Supplementary Table 3), a substantial increase from the approximately 58% coverage of the initial population (23). Alignment to the KitaakeX genome reveals coverage of over 70% of its annotated protein-coding genes (Figure 3-A, Supplementary Table 3). Compared to other major rice mutant resources, such as the *N*-methyl-*N*-nitrosourea (MNU) mutant population has been reported to cover approximately 61% of annotated rice genes (54). Other major rice mutant resources include the POSTECH T-DNA insertion collection (55), the Nagina 22 (N22) EMS mutant resource (56), the IR64 EMS population (57), a genome-scale CRISPR/Cas9 mutant library (58), and Tos17 retrotransposon-tagged lines (59). While several of these collections comprise extensive mutant populations and have contributed significantly to functional studies, most do not report explicit genome-wide gene coverage metrics. In this context, the expanded KitBase population stands out, having comprehensively characterized rice mutant resources currently available, offering high genome coverage.

A novel and technically important aspect of our study was the use of a dual-reference genome alignment strategy. While Nipponbare’s more complete annotation (55,986 genes) facilitated the identification of a larger set of affected genes overall compared to KitaakeX (35,594 genes), leading to ∼8.5% more total mutations detected (Supplementary Table 2). Alignment to the KitaakeX background genome brings specific information for minimizing false negatives due to sequence divergence from Nipponbare (∼0.5% differences) and detecting private alleles unique to Kitaake (47). This approach reduces reference-read mismatches that can mask true mutations. Importantly, both references contributed unique variant calls, with a subset of mutations (∼1–2%) detected exclusively in the KitaakeX alignment (Figure 2-D). The KitBase resource integrates results from both alignment strategies to provide a more comprehensive mutation catalog, and the web-based ID converter tool facilitates navigating between orthologous genes identified by the different annotations (Figure 7-C, Supplementary Figure 3-G).

FN mutagenesis induces a diverse spectrum of genetic alterations, including SBSs, deletions, insertions, inversions, translocations, and tandem duplications (23). In the expanded KitBase population, SBSs and deletions collectively constitute the vast majority (∼80%) of detected variants (Figure 5-B, Supplementary Table 3). Detailed analysis of SBSs reveals a strong predominance of C>T transitions (41.5% of all SBS), with an overall Ti/Tv ratio of 1.44, significantly above the ∼0.5 expected under random mutation and consistent with previous FN studies (16). This elevated transition bias, particularly the C>T signature, is the biochemical hallmark of oxidative base damage and cytosine deamination, suggesting that reactive oxygen species (ROS) generated by ionizing radiation contribute substantially to SBS induction alongside direct double-strand breaks. Notably, TE-associated genes exhibited a significantly higher C>T frequency than non-TE genes (47.9% vs. 37.3%, p = 2.3×10⁻²⁰), consistent with elevated mutation rates at methylated cytosines in transposable element regions (Supplementary Figure 1-L). Deletions are the most prevalent mutation type affecting gene regions and display a bimodal size distribution that provides mechanistic insight into how fast-neutron irradiation damages DNA (Figure 5-D; Supplementary Table 3). Analysis of deletion sizes on a log₁₀ scale revealed two statistically distinct populations separated by a pronounced gap between 51 bp and 10 kb (Figure 5-F). A major peak of micro-deletions ≤50 bp (73.6% of all deletions) generated primarily by NHEJ microresection and replication slippage, and a second peak in the 10 - 100 kb range (19.7%) consistent with DSB induction and imprecise repair by MMEJ. This bimodal pattern was consistent across both alignments (gap containing only 3.8% and 3.2% of deletions respectively). This result was supported by bimodality coefficient analysis (BC = 0.853 and 0.863; threshold 0.555) and Gaussian mixture modeling (ΔBIC > 79,000). These data indicate that the bimodal deletion size distribution is a biological property of FN mutagenesis rather than a reference genome artifact. A similar bimodal deletion spectrum has been reported in fast-neutron-mutagenized *Arabidopsis thaliana* (63), suggesting this is a conserved feature of fast-neutron-induced DNA damage across plant species.

These two deletion size classes serve complementary roles in functional genomics. Small deletions (≤10 bp) provide precise, single-gene knockouts (0.15 genes per deletion on average in Nipponbare) ideal for unambiguous genotype-phenotype association. Large deletions (>100 kb) average 12.6 genes per event but uniquely enable studies of gene redundancy and gene cluster function, as demonstrated in prior KitBase studies identifying structural variants underlying disease resistance and grain morphology phenotypes (47–50).

The analysis of the mutation landscape also provides insights into genome biology. The uniform distribution of mutations across chromosomes suggests the absence of strong mutational hotspots for FN mutagenesis (Figure 4-A and Supplementary Figure 1-C). Furthermore, the majority of unmutated genes have low or no expression and shorter coding regions (approximately 59.6% of unmutated genes have FPKM < 10 and are significantly shorter than expressed genes), which may have been missed by chance (Supplementary Figure 1-F, Supplementary Table 3). The presence of a subset of unmutated genes with moderate to high expression levels (over 3,400 non-TE genes, including 575 highly expressed ones) strongly suggests they represent likely essential genes in rice. Functional enrichment analysis confirms that these highly expressed unmutated genes are significantly over-represented in vital cellular processes such as translation, metabolism, and protein processing (Supplementary Figure 1-G and Supplementary Table 3), aligning with the concept that disruption of essential genes is deleterious and selected against (42,60–62). These genes represent valuable candidates for future functional studies using conditional knockdown or inducible gene-editing strategies.

Additionally, the identification of a substantial number of mutations within intergenic regions (over 169,000 in each alignment) highlights the potential of KitBase for investigating regulatory mutations that can alter gene expression by affecting promoters, enhancers, or other *cis*-regulatory elements (Supplementary Table 1) (63–65).

Another advantage of KitBase mutant collection is the relatively low mutational load per line compared to other mutagens like EMS or gamma rays (Figures 2-B and 4-C), which is consistent with previous findings (16,23). On average, lines carry tens of affected genes (27.09 genes/line in the 90% subset for Nipponbare, 12.2 for KitaakeX). This low background mutation rate increases the precision of genotype-phenotype associations and suggests that only a small segregating population is typically required to identify the causative mutation. Furthermore, the presence of multiple independent mutations within individual genes (over 20,000 genes with two or more hits in Nipponbare) provides valuable allelic variation, enabling fine-scale dissection of gene function and the study of allelic series.

The utility of the KitBase resource has already been demonstrated through its successful application in various research areas, leading to significant gene discoveries. Examples include the identification of genes involved in root behavior (*HK1*), plant architecture, disease resistance (*sxi2/PALD*, *sxi4/DCL2a*, *RBL1*), grain morphology (*gs9-1/BC12*), and fundamental studies on mutation rates and repair mechanisms (47,49,50,52,53,56). These studies showcase KitBase’s power in enabling both traditional forward and reverse genetic approaches, as well as contributing to advanced computational methodologies like training deep learning models for phenotyping (51). Similar gene-trait linkage platforms have emerged in other crops, such as Tnt1-tagged *Medicago truncatula* (66), transposon-insertion lines in tomato (67), FN-induced soybean populations (22), and EMS-induced mutant population in sorghum (65). The ability to identify specific mutations, including structural variants, and link them to diverse phenotypic outcomes underscores KitBase’s value compared to resources with less complete or lower-resolution mutation data.

So far, we have phenotypically characterized hundreds of mutant lines for at least one of the traits related to plant vigor, growth and development, anatomy and morphology, and yield (Figure 6; Supplementary Table 4). The combination of WGS and phenotypic screening enables direct and high-confidence connections between genotype and phenotype, streamlining the discovery of candidate genes (68,69). To ensure maximal accessibility and utility, the updated KitBase web interface (http://kitbase.ucdavis.edu/) serves as a central resource database (Figure 7, Supplementary Figure 3). It provides efficient search functionalities by Line ID, Gene ID, Keyword, or Phenotypic trait, allowing users to quickly locate data of interest. Dedicated pages for each mutant line integrate comprehensive genomic and phenotypic information (Figure 7-E), and the integrated JBrowse genome browser facilitates interactive visualization of all mutations along the chromosomes, aiding in identifying overlapping hits and examining intergenic regions (Supplementary Figure 3-E). Furthermore, the platform offers access to all raw sequencing data (Figure 7-C), downloadable curated data files, essential bioinformatics tools (Figure 7-D, Supplementary Figure 3-F, 3-G), and valuable user support materials, including protocols and webinars. These resources collectively help researchers to effectively leverage the KitBase population for their studies, from initial data exploration to obtaining physical seed stocks via a dedicated ’Seed Order’ page.

In conclusion, the expanded and comprehensively characterized KitBase population, with its high genome coverage, diverse mutation spectrum including structural variants, relatively low mutation load per line, dual-reference alignment data, integrated genomic and phenotypic information, and user-friendly web interface, represents a pivotal resource for functional genomics research in rice. Its proven utility in identifying genes for various agronomic traits, enabling advanced phenotyping approaches, and contributing to fundamental biological insights positions KitBase as a cornerstone platform to help our understanding of gene function and trait development in this vital crop.

## Data availability

The database described in this article is freely available online at https://kitbase.ucdavis.edu/.

## Acknowledgements

We are deeply grateful to Maria E. Hernandez for her valuable contributions to this project. We also thank the students and colleagues whose names are not individually listed here for their dedicated support with genomic DNA isolation, sample submission, seed organization, and data processing. Their advice, suggestions, and assistance greatly strengthened the quality of this work, and we sincerely appreciate their commitment.

---

**Supplementary Figure 1- Overview of Mutation Distribution, Gene Impact, and Functional Classification in the KitaakeX FN-Mutagenized Rice Population. (A)** Histogram representing the overall mutation frequency distribution per line in both alignments. The analysis confirms the normality of the distribution and supports the consistency of mutation-detection patterns. (**B)** Venn Diagram showing the similarity between the affected genes found in the Nipponbare alignment and the identified putative orthologs affected genes from the KitaakeX alignment. In black, the total annotated genes, and in blue, the non-TE genes. **(C)** Chromosomal mapping and distribution of affected genes in the KitaakeX alignment. The heatmap represents the number of distinct mutations per gene, ranging from yellow (one mutation) to red (ten or more mutations). Non-mutated genes are shown in black, and intergenic regions are depicted in white. **(D)** Venn diagram illustrating the overlap between the transcription factor genes identified in the PlantTFDB database and those affected by FN-induced mutations in the KitBase. This visualization highlights the extent of the mutational impact on the transcription factor repertoire within the Nipponbare alignment. **(E)** Comparison of the total number of genes per transcription factor (TF) family (gray bars) versus the number of affected genes within each family (marine blue bars). This analysis highlights the differential impact of mutations across various TF families. **(F)** Heatmap plot of the *in silico* expression analysis for non-affected genes in the Kitaake rice mutant population, aligned to the Nipponbare reference genome. Expression data were collected across 13 tissue types using 682 RNA-seq datasets from the Rice RNA-seq Database (https://plantrnadb.com/ricerna/). **(G-H)** Gene Ontology (GO) enrichment analysis of **(G)** Biological Process and **(H)** Kegg for non-affected genes with high expression levels (based on *in silico* analysis) in the Kitaake rice mutant population, aligned to the Nipponbare reference genome. **(I)** Distribution of gene lengths among non-mutated genes, categorized by expression levels. Expression data and genes were classified into five categories: unidentified, noise, low, medium, and high expression. Gene length information was sourced from the MSU database (Kawahara *et al.*, 2013). (**J)** Chromosomal Distribution of Non-mutated genes. In red are the non-TE genes with high *in silic* expression; grey are the other non-TE genes and TE genes. **(L)** SBS Spectrum in Gene Regions Stratified by Gene Classification. SBS6 frequency distribution for single-base substitutions occurring within gene bodies, stratified by gene classification: Nipponbare non-TE genes, Nipponbare TE-related genes, KitaakeX genes with confident Nipponbare ortholog assignments classified as non-TE, and KitaakeX genes lacking a confident ortholog assignment (Unmapped). Statistical significance of the difference in SBS6 spectrum between Nipponbare non-TE and TE genes was assessed using a chi-square test of independence (χ² = 101.7, p = 2.3×10⁻²⁰). KitaakeX TE gene classification via ortholog mapping yielded insufficient sample size for reliable spectrum estimation and is not shown; this likely reflects the low ortholog conservation of transposable element sequences between cultivars. **(M)** Ti/Tv ratios for each gene category, with the genome-wide Ti/Tv (1.44) shown as a dashed reference line.

**Supplementary Figure 2. Phenotypic variation, trait correlations, and multivariate analysis of KitaakeX FN-mutant rice lines. (A)** Distribution of phenotypic variation across all evaluated traits in the FN-mutant population, including germination rate (seedling vigor), albino seedling frequency, tiller number, days to heading (flowering time), plant height, panicle traits (length, weight, and filled grain number), and seed traits (total seed number per panicle and fertility). Trait values were normalized relative to the wild-type control (KitaakeX), where a value of 1 (100%) represents wild-type performance, allowing comparison across planting batches. (**B)** Correlation matrix of all phenotypic traits. Asterisks (*) indicate statistically significant correlations (p < 0.05). Color intensity represents the strength of correlation coefficients, ranging from blue (low correlation) to red (high correlation). **(C)** Principal Component Analysis (PCA) separating mutant lines based on phenotypic variation. The analysis highlights clustering by plant height and panicle weight. Color-coded groups include: blue (dwarf stature with low panicle weight), green (wild-type height with low panicle weight), and red (wild-type-like in both height and panicle weight).

**Supplementary Figure 3 Navigation and Tools in KitBase. (A)** Mutant ID Search Results: Displays the detailed information page for a specific mutant line, including all associated data. The genetic information presented is based on the Nipponbare alignment. **(B)** Gene ID Search Results: Lists all mutant lines with mutations in the specified gene. The first column indicates the mutant line name, followed by columns detailing the genomic information of each mutation. **(C)** Keyword Search Results: Provides a list of mutant lines associated with genes containing the specified keyword (e.g., ’iron’) in their descriptions. **(D)** Phenotype Search Results: Displays available phenotype data for selection. Upon choosing a specific phenotype, a list of characterized mutant lines is presented. **(E)** JBrowse Genome Viewer: Snapshot illustrating mutations within a genomic region across the mutant population. (**F)** BLAST Tool Interface: Page showcasing the BLAST functionality within KitBase. (**G)** ID Converter Tool: Interface for converting between Kitaake and Nipponbare gene IDs, and vice versa. **(H)** Seed Order Form: Page displaying the form required to request seed samples.

**Supplementary Figure 4. KitBase-enabled identification and allelic validation of** *D1/RGA1* **dwarf alleles. (A)** Gene structure of *D1/RGA1* (LOC_Os05g26890) based on the Nipponbare reference genome. Gray boxes indicate exons; lines indicate introns. FN3664-S carries a 136-kb deletion on chromosome 5 (15,481,001-15,617,000 bp) that removes the entire *D1/RGA1* locus. FN1535-S carries a chromosome 5 inversion whose breakpoint falls within exon 4 (marked with a black cross symbol). *D1/RGA1* is the only gene mutated in both lines.**(B)** Representative PCR gel confirming segregation of the *D1/RGA1* deletion in the FN3664-S population. Primers flanking the deletion produce a 979 bp band from the wild-type allele; absence of the band indicates homozygosity for the mutation. Sanger sequencing of the PCR product confirmed that primers amplify the expected genomic region. The same primer set was used to confirm the presence or absence of the mutation in FN1535-S, where a band indicates the presence of at least one wild-type allele. **(C)** Growth curves showing mean plant height (± SD) for KitaakeX, FN3664-S WT-sibling (FN3664-S WT), FN3664-S Dwarf, and FN1535-S at Days 10, 17, 20, 24, 26, 29, 33, 37, and 40 after germination. **(D-G)** Boxplots showing **(D)** panicle length (cm), **(E)** panicle area (cm²), **(F)** seed length (cm), and **(G)** seed area (cm²) for KitaakeX, FN3664-S WT, FN3664-S Dwarf, and FN1535-S. Each point represents one biological replicate. Statistical analysis was performed using one-way ANOVA followed by Tukey’s HSD test; different letters indicate significant differences between groups (p < 0.05). **(H-J)** Representative photographs of KitaakeX, FN3664-S WT-sibling, FN3664-S Dwarf, and FN1535-S showing **(H)** full plant at 40 days after germination, **(I)** panicle, and **(J)** seeds. Scale bar = 1 cm.

**Supplementary Table 1. Comprehensive summary of genome sequencing data for fast-neutron (FN) mutant rice lines analyzed in this study using dual-reference alignment against the Nipponbare and KitaakeX genomes.** The first set of tables provides a comprehensive comparison of the KitaakeX and Nipponbare reference genome assemblies and annotations, organized into four sections. The second set of table details, for each mutant line, the number of mutations and affected genes identified for both alignments. Additionally, it includes information on each mutant ID, generation, and seed availability.

**Supplementary Table 2. Comprehensive Catalog of Mutations Identified in the Fast-Neutron-Induced Kitaake Rice Mutant Population.** This file provides a detailed inventory of the mutations detected across the fast-neutron-induced Kitaake rice mutant population, with mutations identified through alignment to the Nipponbare and KitaakeX reference genome.

**Supplementary Table 3. A comprehensive summary of affected genes in the Kitaake rice mutant population with the Nipponbare and KitaakeX alignments**.

**Supplementary Table 4. Definitions of all data fields for phenotypic measurements recorded in the mutant rice lines analyzed in this study.** Quantitative traits include germination rate, frequency of albino plantlets, tiller number, days to heading, filled grain number, total seed number, seed yield, panicle weight, and plant height. The table also lists qualitative phenotypic observations recorded across the mutant population.

**Supplementary Table 5. KitBase-assisted identification of FN mutant lines with dwarf or semi-dwarf phenotype and candidate causal genes.** Set 1: FN lines carrying mutations in genes previously associated with dwarf or semi-dwarf phenotype in rice, identified through KitBase reverse-genetic search. Columns show the candidate gene name, gene ID (LOC identifier), and the FN lines harboring mutations in that gene. Set 2: Complete mutation profiles of FN1535-S and FN3664-S from KitBase, based on alignment to both the Nipponbare and KitaakeX reference genomes. For each line, the mutation type, chromosome, start position, end position, and affected gene are listed.

