## Supplementary Figures for "Expanding KitBase: Genome and Phenotype Integration of 3,268 Fast-Neutron Rice Mutants"

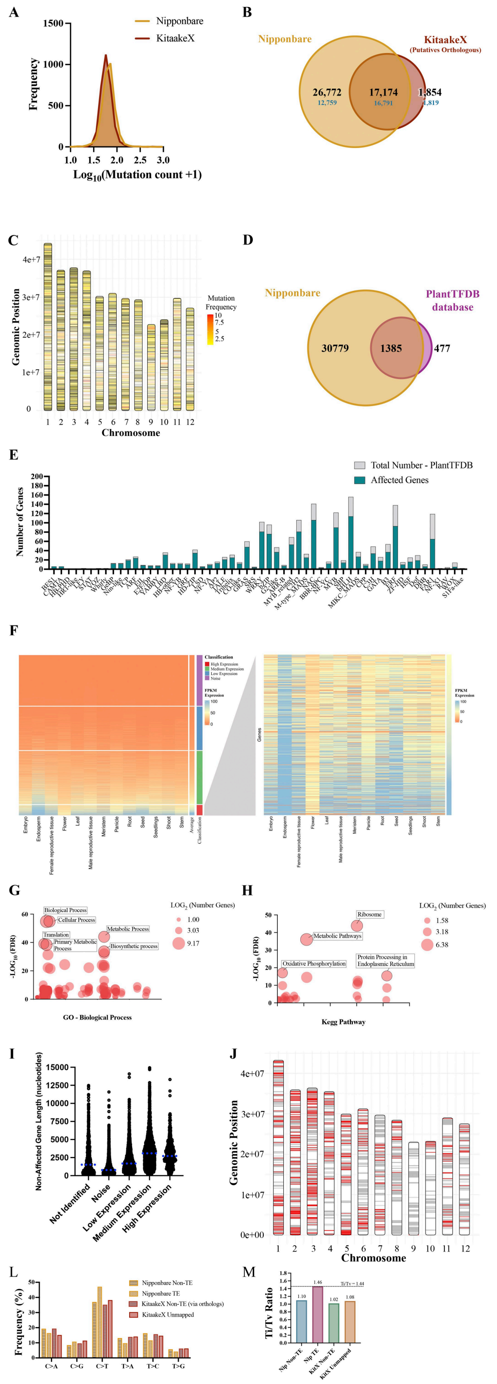


**Supplementary Figure 1- Overview of Mutation Distribution, Gene Impact, and Functional Classification in the KitaakeX FN-Mutagenized Rice Population. (A)** Histogram representing the overall mutation frequency distribution per line in both alignments. The analysis confirms the normality of the distribution and supports the consistency of mutation-detection patterns. **(B)** Venn Diagram showing the similarity between the affected genes found in the Nipponbare alignment and the identified putative orthologs affected genes from the KitaakeX alignment. In black, the total annotated genes, and in blue, the non-TE genes. **(C)** Chromosomal mapping and distribution of affected genes in the KitaakeX alignment. The heatmap represents the number of distinct mutations per gene, ranging from yellow (one mutation) to red (ten or more mutations). Non-mutated genes are shown in black, and intergenic regions are depicted in white. **(D)** Venn diagram illustrating the overlap between the transcription factor genes identified in the PlantTFDB database and those affected by FN-induced mutations in the KitBase. This visualization highlights the extent of the mutational impact on the transcription factor repertoire within the Nipponbare alignment. **(E)** Comparison of the total number of genes per transcription factor (TF) family (gray bars) versus the number of affected genes within each family (marine blue bars). This analysis highlights the differential impact of mutations across various TF families. **(F)** Heatmap plot of the *in silico* expression analysis for non-affected genes in the Kitaake rice mutant population, aligned to the Nipponbare reference genome. Expression data were collected across 13 tissue types using 682 RNA-seq datasets from the Rice RNA-seq Database (https://plantrnadb.com/ricerna/). **(G-H)** Gene Ontology (GO) enrichment analysis of **(G)** Biological Process and **(H)** Kegg for non-affected genes with high expression levels (based on *in silico* analysis) in the Kitaake rice mutant population, aligned to the Nipponbare reference genome. **(I)** Distribution of gene lengths among non-mutated genes, categorized by expression levels. Expression data and genes were classified into five categories: unidentified, noise, low, medium, and high expression. Gene length information was sourced from the MSU database (Kawahara *et al.*, 2013). **(J)** Chromosomal Distribution of Non-mutated genes. In red are the non-TE genes with high *in silic* expression; grey are the other non-TE genes and TE genes. **(L)** SBS Spectrum in Gene Regions Stratified by Gene Classification. SBS6 frequency distribution for single-base substitutions occurring within gene bodies, stratified by gene classification: Nipponbare non-TE genes, Nipponbare TE-related genes, KitaakeX genes with confident Nipponbare ortholog assignments classified as non-TE, and KitaakeX genes lacking a confident ortholog assignment (Unmapped). Statistical significance of the difference in SBS6 spectrum between Nipponbare non-TE and TE genes was assessed using a chi-square test of independence ($\chi^2 = 101.7$, $p = 2.3\times10^{-20}$). KitaakeX TE gene classification via ortholog mapping yielded insufficient sample size for reliable spectrum estimation and is not shown; this likely reflects the low ortholog conservation of transposable element sequences between cultivars. **(M)** Ti/Tv ratios for each gene category, with the genome-wide Ti/Tv (1.44) shown as a dashed reference line.

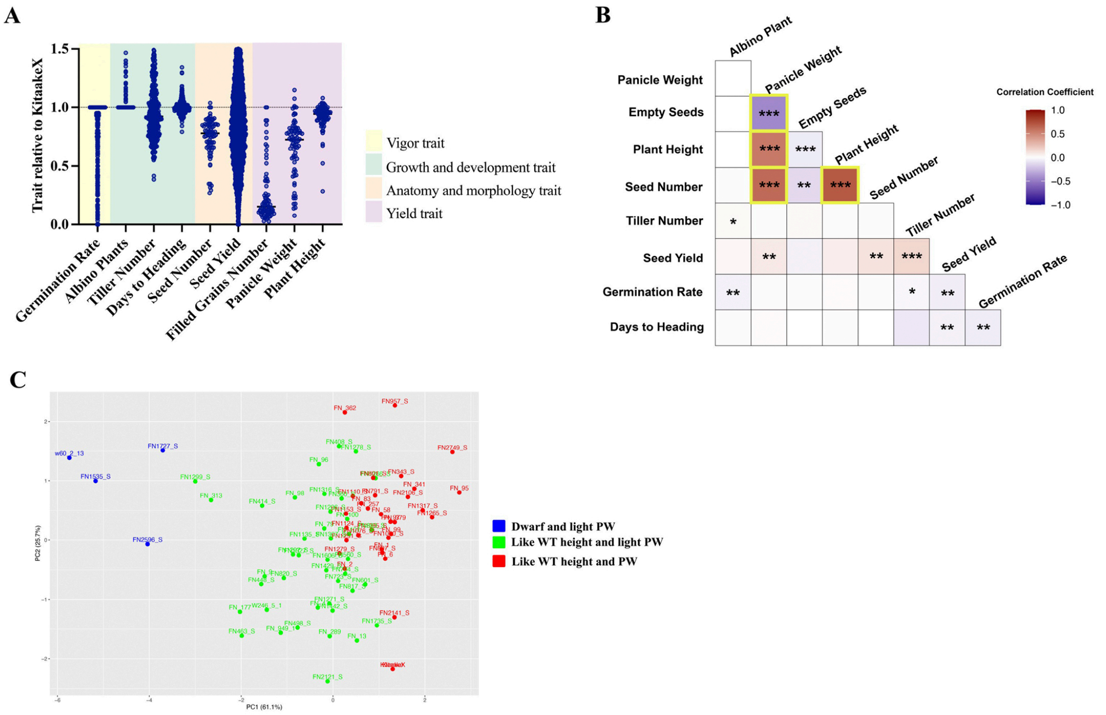


**Supplementary Figure 2. Phenotypic variation, trait correlations, and multivariate analysis of KitaakeX FN-mutant rice lines. (A)** Distribution of phenotypic variation across all evaluated traits in the FN-mutant population, including germination rate (seedling vigor), albino seedling frequency, tiller number, days to heading (flowering time), plant height, panicle traits (length, weight, and filled grain number), and seed traits (total seed number per panicle and fertility). Trait values were normalized relative to the wild-type control (KitaakeX), where a value of 1 (100%) represents wild-type performance, allowing comparison across planting batches. (**B)** Correlation matrix of all phenotypic traits. Asterisks (*) indicate statistically significant correlations ($p < 0.05$). Color intensity represents the strength of correlation coefficients, ranging from blue (low correlation) to red (high correlation). **(C)** Principal Component Analysis (PCA) separates mutant lines based on phenotypic variation. The analysis highlights clustering by plant height and panicle weight. Color-coded groups include: blue (dwarf stature with low panicle weight), green (wild-type height with low panicle weight), and red (wild-type-like in both height and panicle weight).

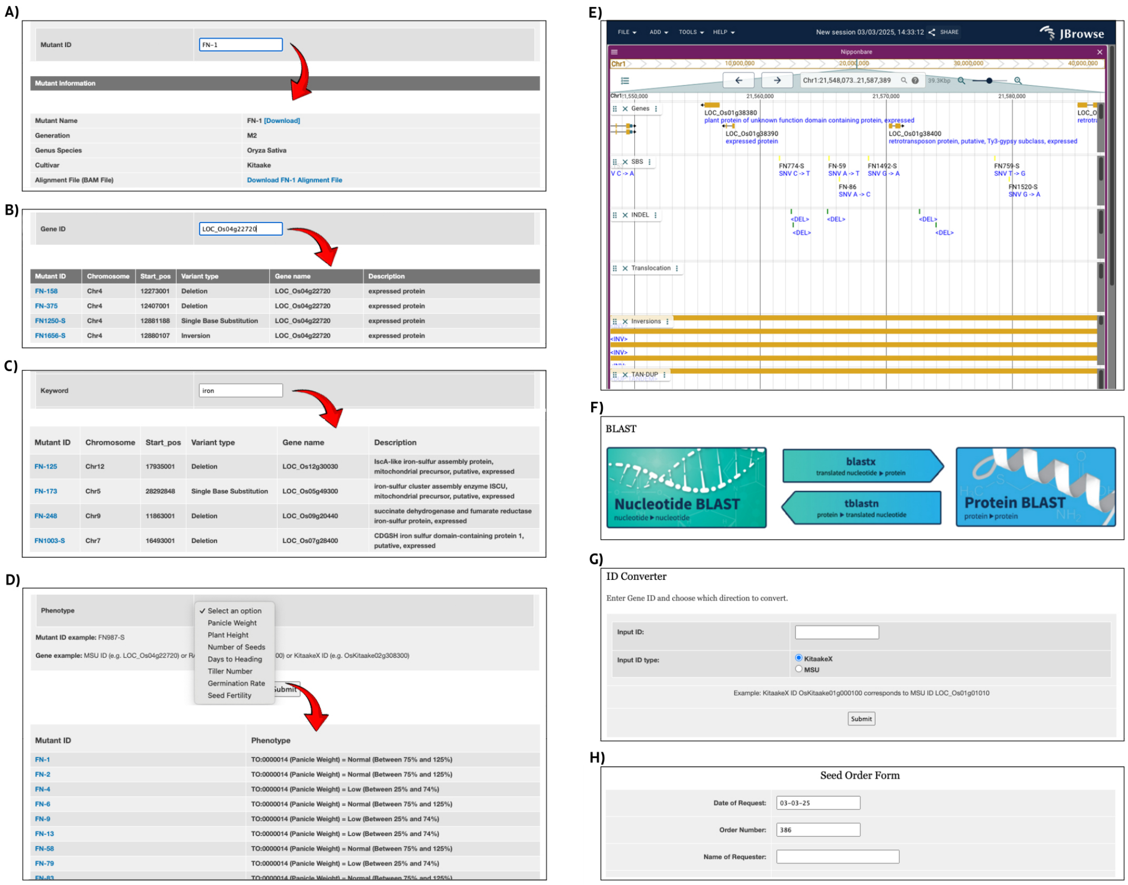

**Supplementary Figure 3 Navigation and Tools in KitBase. (A)** Mutant ID Search Results: Displays the detailed information page for a specific mutant line, including all associated data. The genetic information presented is based on the Nipponbare alignment. **(B)** Gene ID Search Results: Lists all mutant lines with mutations in the specified gene. The first column indicates the mutant line name, followed by columns detailing the genomic information of each mutation. **(C)** Keyword Search Results: Provides a list of mutant lines associated with genes containing the specified keyword (e.g., 'iron') in their descriptions. **(D)** Phenotype Search Results: Displays available phenotype data for selection. Upon choosing a specific phenotype, a list of characterized mutant lines is presented. **(E)** JBrowse Genome Viewer: Snapshot illustrating mutations within a genomic region across the mutant population. **(F)** BLAST Tool Interface: Page showcasing the BLAST functionality within KitBase. **(G)** ID Converter Tool: Interface for converting between Kitaake and Nipponbare gene IDs, and vice versa. **(H)** Seed Order Form: Page displaying the form required to request seed samples.

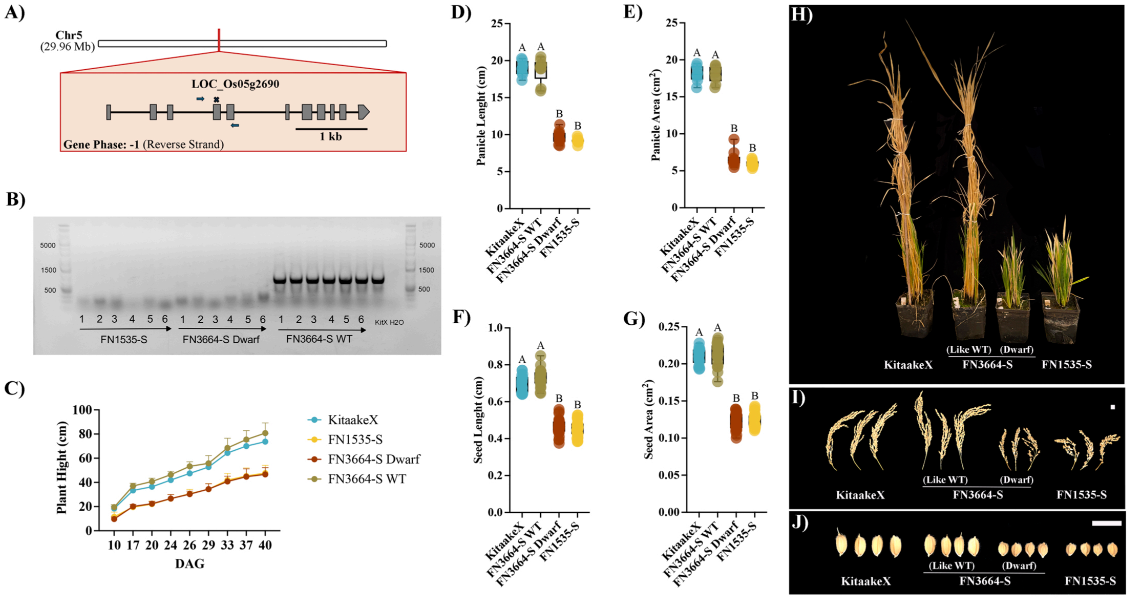


**Supplementary Figure 4. KitBase-enabled identification and allelic validation of *D1/RGA1* dwarf alleles.** **(A)** Gene structure of *D1/RGA1* (LOC_Os05g26890) based on the Nipponbare reference genome. Gray boxes indicate exons; lines indicate introns. FN3664-S carries a 136-kb deletion on chromosome 5 (15,481,001-15,617,000 bp) that removes the entire *D1/RGA1* locus. FN1535-S carries a chromosome 5 inversion whose breakpoint falls within exon 4 (marked with a black cross symbol). *D1/RGA1* is the only gene mutated in both lines. **(B)** Representative PCR gel confirming segregation of the *D1/RGA1* deletion in the FN3664-S population. Primers flanking the deletion produce a 979 bp band from the wild-type allele; absence of the band indicates homozygosity for the mutation. Sanger sequencing of the PCR product confirmed that primers amplify the expected genomic region. The same primer set was used to confirm the presence or absence of the mutation in FN1535-S, where a band indicates the presence of at least one wild-type allele. **(C)** Growth curves showing mean plant height (± SD) for KitaakeX, FN3664-S WT-sibling (FN3664-S WT), FN3664-S Dwarf, and FN1535-S at Days 10, 17, 20, 24, 26, 29, 33, 37, and 40 after germination. **(D-G)** Boxplots showing **(D)** panicle length (cm), **(E)** panicle area ($cm^2$), **(F)** seed length (cm), and **(G)** seed area ($cm^2$) for KitaakeX, FN3664-S WT, FN3664-S Dwarf, and FN1535-S. Each point represents one biological replicate. Statistical analysis was performed using one-way ANOVA followed by Tukey's HSD test; different letters indicate significant differences between groups ($p < 0.05$). **(H-J)** Representative photographs of KitaakeX, FN3664-S WT-sibling, FN3664-S Dwarf, and FN1535-S showing **(H)** full plant at 40 days after germination, **(I)** panicle, and **(J)** seeds. Scale bar = 1 cm.
